# Recombination patterns in coronaviruses

**DOI:** 10.1101/2021.04.28.441806

**Authors:** Nicola F. Müller, Kathryn E. Kistler, Trevor Bedford

## Abstract

As shown during the SARS-CoV-2 pandemic, phylogenetic and phylodynamic methods are essential tools to study the spread and evolution of pathogens. One of the central assumptions of these methods is that the shared history of pathogens isolated from different hosts can be described by a branching phylogenetic tree. Recombination breaks this assumption. This makes it problematic to apply phylogenetic methods to study recombining pathogens, including, for example, coronaviruses. Here, we introduce a Markov chain Monte Carlo approach that allows inference of recombination networks from genetic sequence data under a template switching model of recombination. Using this method, we first show that recombination is extremely common in the evolutionary history of SARS-like coronaviruses. We then show how recombination rates across the genome of the human seasonal coronaviruses 229E, OC43 and NL63 vary with rates of adaptation. This suggests that recombination could be beneficial to fitness of human seasonal coronaviruses. Additionally, this work sets the stage for Bayesian phylogenetic tracking of the spread and evolution of SARS-CoV-2 in the future, even as recombinant viruses become prevalent.

## Main

Since its emergence, genetic sequence data has been applied to study the evolution and spread of SARS-CoV-2. Genetic sequences have, for example, been used to investigate natural versus lab origins of SARS-CoV-2 (Andersen *et al*., 2020), when SARS-CoV-2 was introduced into the US (Bedford *et al*., 2020) as well as whether genetic variants differ in growth rate (Volz *et al*., 2021). These analyses often rely on phylogenetic and phylodynamic approaches, at the heart of which are phylogenetic trees. Such trees denote how viruses isolated from different individuals are related and contain information about the transmission dynamics connecting these infections (Grenfell *et al*., 2004).

Along with mutations introduced by errors during replication or by anti-viral molecules (for example (Kim *et al*., 2014)), different recombination processes contribute to genetic diversity in RNA viruses (reviewed by Simon-Loriere and Holmes, 2011). Reassortment in segmented viruses (generally negative-sense RNA viruses), such as influenza or rotaviruses, can produce offspring that carry segments from different parent lineages (McDonald *et al*., 2016). In other RNA viruses (generally positive-sense RNA viruses), such as flaviviruses and coronaviruses, homologous recombination can combine different parts of a genome from different parent lineages in absence of physically separate segments on the genome of those viruses (Su *et al*., 2016). The main mechanism of this process is thought to be via template switching (Lai, 1992), where the template for replication is switched during the replication process. Recombination breakpoints in experiments appear to be largely random, with selection selecting recombination breakpoints in some areas of the genome (Banner and Mc Lai, 1991). Recent work shows that recombination breakpoints occur more frequently in the spike region of beta-coronaviruses, such as SARS-CoV-2 (Bobay *et al*., 2020). While the evolutionary purpose of recombination in RNA viruses is not completely understood (Simon-Loriere and Holmes, 2011), there are different explanations of why recombination may be beneficial. In general, recombination is selected for if breaking up the linkage disequilibrium is beneficial (Barton, 1995). Recombination can help purge deleterious mutations from the genome, such as proposed by the mutational-deterministic hypothesis (Feldman *et al*., 1980). It can also increase the rate at which fit combination of mutations occur, such as stated by the Robertson-Hill effect (Hill and Robertson, 1966). Alternatively, recombination in RNA viruses may also just be a by-product of the processivity the viral polymerase (Simon-Loriere and Holmes, 2011).

Recombination poses a unique challenge to phylogenetic methods, as it violates the very central assumption that the evolutionary history of individuals can be denoted by branching phylogenetic trees. Recombination breaks this assumption and requires representation of the shared ancestry of a set of sequences as a network. Not accounting for this can lead to biased phylogenetic and phylodynamic inferences (Posada and Crandall, 2002; Müller *et al*., 2020). An analytic description of recombination is provided by the coalescent with recombination, which models a backwards in time process where lineages can coalesce and recombine (Hudson, 1983). When recombination is considered backwards in time, a single lineage results in two parent lineages, with one parent lineage carrying the genetic material from one side of a random recombination breakpoint and the other parent lineage carrying the genetic material of the other side of this breakpoint. This equates to the backwards in time equivalent of template switching where there is one recombination breakpoint per recombination event.

Currently, some Bayesian phylogenetic approaches exist that infer recombination networks, or ancestral recombination graphs (ARG), but are either approximate or do not directly allow for efficient model-based inference. Some approaches consider tree-based networks (Didelot *et al*., 2010; Vaughan *et al*., 2017), where the networks consist of a base tree with recombination edges that always attach to edges on the base tree. Alternative approaches rely on approximations to the coalescent with recombination (Rasmussen *et al*., 2014; McVean and Cardin, 2005), consider a different model of recombination (Müller *et al*., 2020), or seek to infer recombination networks absent an explicit recombination model (Bloomquist and Suchard, 2010). Bayesian and maximum likelihood methods have also been used to account for gene transfer events when describing the evolutionary history of species from multiple loci (for example (Meng and Kubatko, 2009; Yu *et al*., 2014). Additionally, methods have been used to describe non-tree-like evolution using split trees (Bryant and Moulton, 2004; Huson and Bryant, 2006). There is, however, a gap for Bayesian inference of recombination networks under the coalescent with recombination that can be applied to study pathogens, such as coronaviruses.

In order to fill this gap, we here develop a Markov chain Monte Carlo (MCMC) approach to efficiently infer recombination networks under the coalescent with recombination for sequences sampled over time. This framework allows joint estimation of recombination networks, effective population sizes, recombination rates and parameters describing mutations over time from genetic sequence data sampled through time. We explicitly do not make additional approximation to characterize the recombination process, other than those of the coalescent with recombination (Hudson, 1983), such as, for example, the approximation of tree based networks. We implemented this approach as an open source software package for BEAST2 (Bouckaert *et al*., 2018). This allows incorporation of the various evolutionary models already implemented in BEAST2. Using a Bayesian approach has several advantages. In particular, it allows us to account for uncertainty in the parameter and network estimates. Additionally, it allows balancing different sources of information against each other. The coalescent with recombination model, for example, will tend to favor networks with fewer recombination events. This “cost” of adding more recombination events depends on the recombination rate. At lower rates of recombination, adding new recombination events is more costly and the information coming from the sequence alignment in support of a recombination event needs to be greater.

We first apply the coalescent with recombination to study the evolutionary history of SARS-like coronaviruses. Doing so, we show that the evolutionary history of SARS-like coronaviruses is extremely complex and has little resemblance to tree-like evolution. Additionally, we show that recombination only occurred between closely related SARS-like viruses in the recent history of SARS-CoV-2. Next, we reconstruct the evolutionary histories of MERS-CoV and three seasonal human coronaviruses to show that recombination also frequently occurs in human coronaviruses at rates that are comparable to reassortment rates in influenza viruses. Next, we show that recombination breakpoints in human coronaviruses vary with rates of adaptation across the genomes, suggesting that recombination events are positively or negatively selected based on where breakpoints occur.

### Rampant recombination in SARS-like coronaviruses

Recombination has been implicated at the beginning of the SARS-CoV-1 outbreak (Hon *et al*., 2008) and has been suggested as the origin of the receptor binding domain in SARS-CoV-2 (Li *et al*., 2020), though Boni *et al*. (2020) report that recombination is unlikely to be the origin of SARS-CoV-2. While this strongly suggests non-tree-like evolution, the evolutionary history of SARS-like viruses has, out of necessity, mainly been denoted using phylogenetic trees.

We here reconstruct the recombination history of SARS-like viruses, which includes SARS-CoV-1 and SARS-CoV-2 as well as related bat (Ge *et al*., 2013, 2016; Zhou *et al*., 2020) and pangolin (Lam *et al*., 2020) coronaviruses. To do so, we infer the recombination network of SARS-like viruses under the coalescent with recombination. We assumed that the rates of recombination and effective population sizes were constant over time and that the genomes evolved under a GTR+Γ_4_ model. Similar to the estimate in Boni *et al*. (2020), we used a fixed evolutionary rate of 5 × 10^−4^ per nucleotide and year. We fixed the evolutionary rate, since the time interval of sampling between individual isolates is relatively short compared to the time scale of the evolutionary history of SARS-like viruses. This means that the sampling times themselves offer little insight into the evolutionary rates and, in absence of other calibration points, there is little information about the evolutionary rate in this dataset. This, in turn, means that if the evolutionary rate we used here is inaccurate then the timings of common ancestors will also be inaccurate. Therefore, exact timings and calendar dates in this analyses should be taken as guide posts rather than formal estimates.

As shown in Figure 1A, the evolutionary history of SARS-like viruses is characterized by frequent recombination events. Consequently, characterizing evolutionary history of SARS-like viruses by a single genome-wide phylogeny is bound to be inaccurate and potentially misleading. We infer the recombination rate in SARS-like viruses to be approximately 2 × 10^−6^ per site per year, which is about 200 times lower than the evolutionary rate. This rate translated to about 0.06 recombination events per lineage per year, which is slightly lower than the estimated rate of recombination for the seasonal human coronaviruses and the reassortment rates for pandemic 1918 like influenza A/H1N1 and influenza B viruses, which are all around 0.1 – 0.2 reassortment events per lineage per year (Müller *et al*., 2020). This recombination rate is a function of co-infection rates, probability of recombination occurring upon co-infection, and selection. As such, the recombination rate we infer here will be (possibly substantially) lower than the within host rate of recombination.

**Figure 1:**
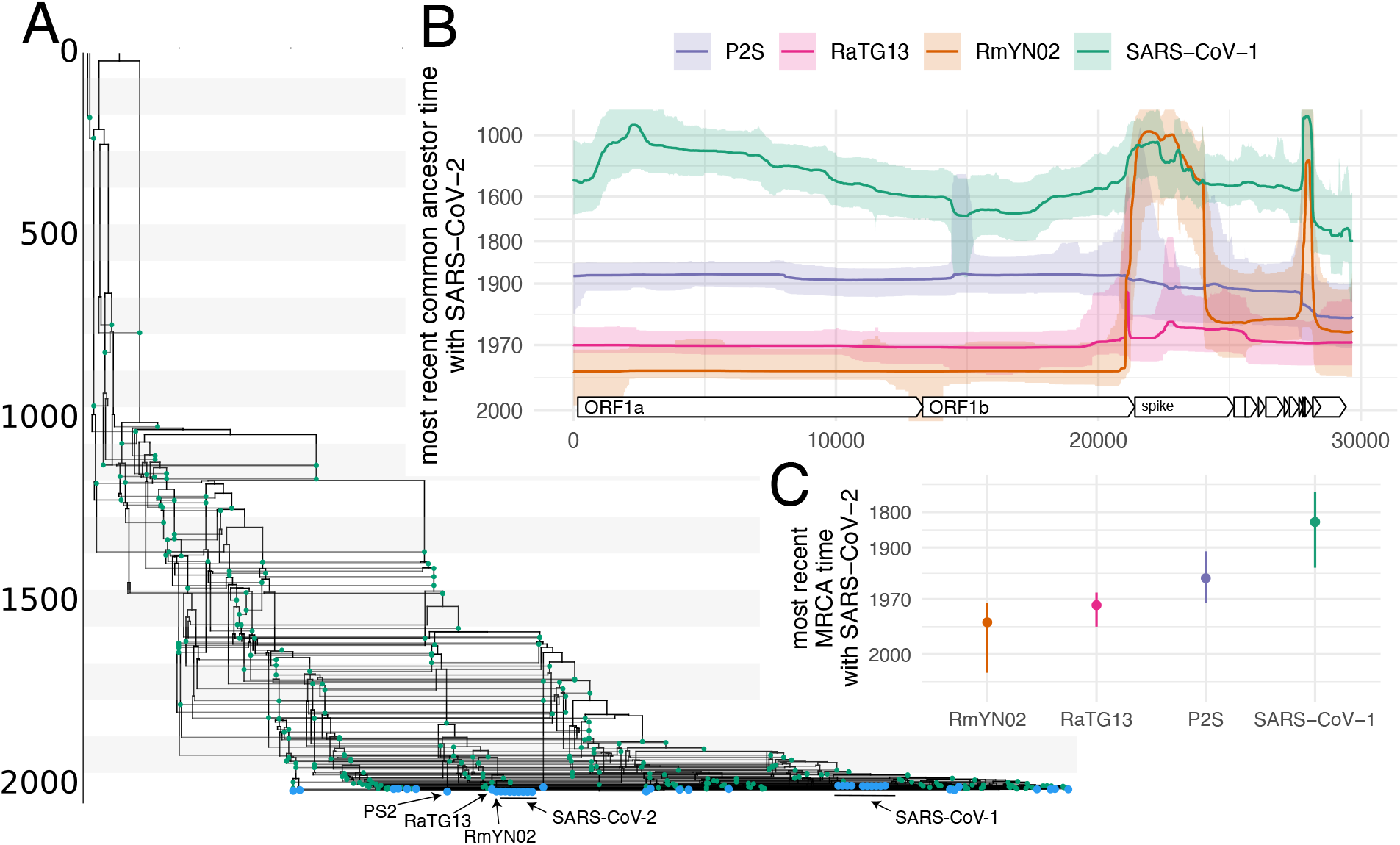
Evolutionary history of SARS-like viruses. **A** Maximum clade credibility network of SARS-like viruses. Blue dots denote samples and green dots recombination events. **B** Common ancestor times of Wuhan-Hu1 (SARS-CoV-2) with different SARS-like viruses on different positions of the genome. The y-axis denotes common ancestor times in log scale. **C** Most recent time anywhere on the genome that Wuhan-Hu1 shared a common ancestor with different SARS-like viruses

These recombination events were not evenly distributed across the genome and, instead, were relatively higher in areas outside those coding for ORF1ab (Fig. S1 & S5). Additionally, our inference suggests that rates of recombination are slightly elevated on spike subunit 1 compared to subunit 2 (Fig. S1). If we track recombination events ancestral to the SARS-CoV-2 lineage that are inferred to have happened in the last 100 years, we find evidence for recombination breakpoints occurring close to the 5’ end of spike, just outside the coding region (see fig S4). Additionally, we find support for recombination breakpoints toward the 3’ end of spike, near the nucleocapsid gene (see fig S4). If we assume that during genome replication in coronaviruses template shifts occur randomly on the genome (Banner and Mc Lai, 1991), differences in observed recombination rates could be explained by selection favoring recombination events 3’ to ORF1ab relative to elsewhere on the genome. For instance, it has been suggested that selection has acted on multiple recombination events within spike to enhance dynamic molecular movements of the Spike protein (Tagliamonte *et al*., 2021).

We next investigate when different viruses last shared a common ancestor (MRCA) along the genome (see Fig. 1B and Fig. S2). RmYN02 (Zhou *et al*., 2020) shares the MRCA with SARS-CoV-2 on the part of the genome that codes for ORF1ab (Fig. 1B). We additionally find strong evidence for one or more recombination events in the ancestry of RmYN02 at the beginning of spike (Fig. 1B). This recent recombination event is unlikely to have occurred with a recent ancestor of any of the coronaviruses included in this dataset since the common ancestor of RmYN02 with any other virus in the dataset is approximately the same (Fig. S3A). In other words, large parts of the spike protein of RmYN02 are as related to SARS-CoV-2 as SARS-CoV-2 is to SARS-CoV-1. The common ancestor timings of P2S across the genome are equal between RaTG13 and SARS-CoV-2 (Fig. S3C). RaTG13 on the other hand is more closely related to SARS-CoV-2 than P2S (Fig. S3B) across the entire genome.

When looking at when different viruses last shared a common ancestor anywhere on the genome (in other words: when the ancestral lineages of two viruses last crossed paths), we find that RmYN02 has the most recent MRCA with SARS-CoV-2 (Fig. S3C). The median estimate of the most recent MRCA between SARS-CoV-2 and RmYN02 is 1986 (95% CI: 1973–2005). For RaTG13 it is 1975 (95% CI: 1988–1964), for P2S it is 1949 (95% CI: 1907–1973) and with SARS-CoV-1 it is 1834 (95% CI: 1707–1935). These estimates are contingent on a fixed evolutionary rate of 5 × 10^−4^ per nucleotide per year.

### Rates of recombination vary with rates of adaptation in human seasonal coronaviruses

We next investigate recombination patterns in MERS-CoV, which has over 2500 confirmed cases in humans, as well as in human seasonal coronaviruses 229E, OC43 and NL63, which have widespread seasonal circulation in humans. As for the SARS-like viruses, we jointly infer recombination networks, rates of recombination and population sizes for these viruses. We assumed that the genomes evolved under a GTR+Γ_4_ model and, in contrast to the analysis of SARS-like viruses, inferred the evolutionary rates. We observe frequent recombination in the history of all 4 viruses, wherein genetic ancestry is described by network rather than a strictly branching phylogeny (Fig. 2A-D).

**Figure 2:**
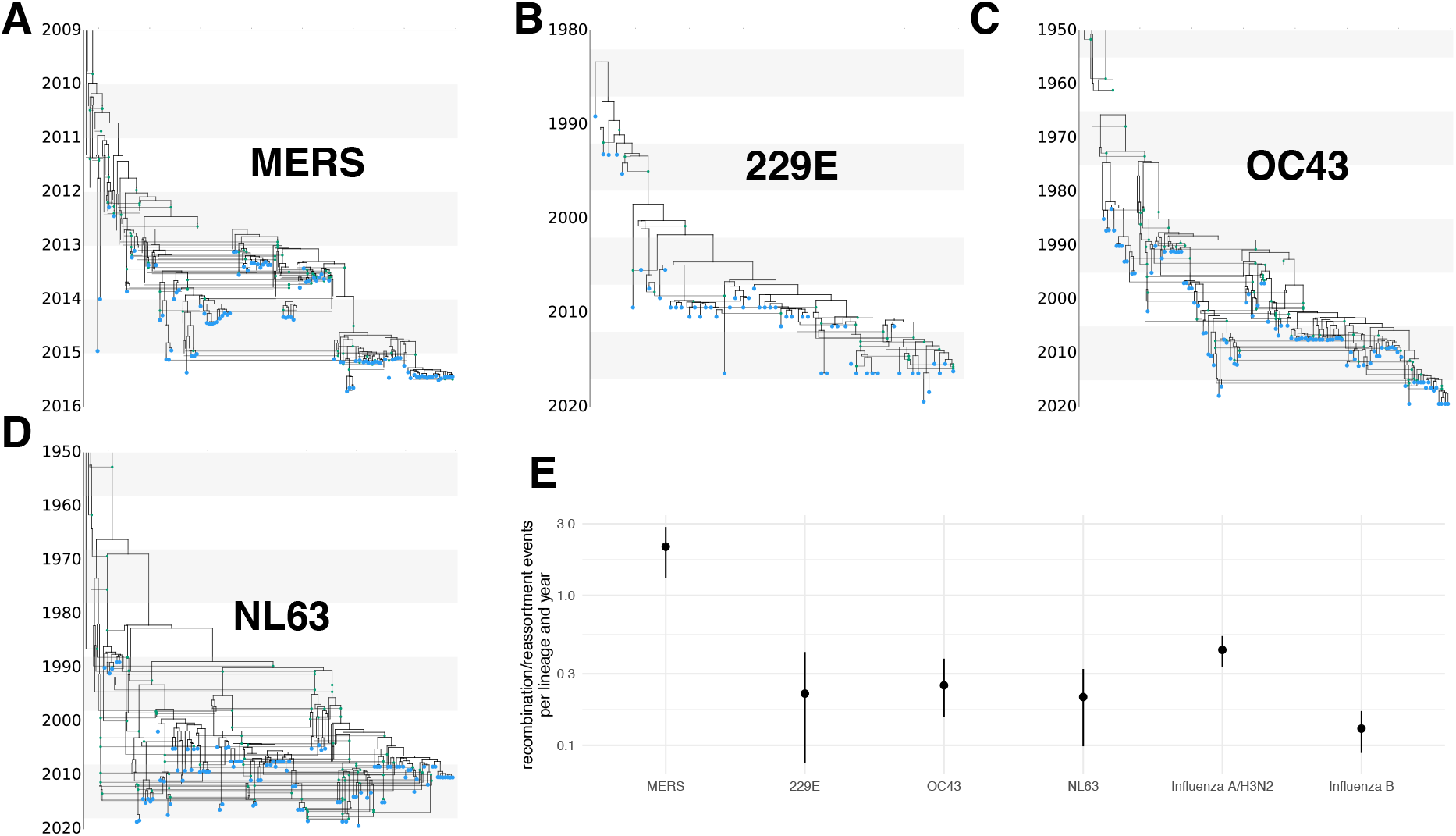
Recombination networks and rates for coronaviruses MERS, 229E, OC43 and NL63. Recombination networks for MERS (**A**) and seasonal human coronaviruses 229E (**B**), OC43 (**C**) and NL63 (**D**). **E** Recombination rates (per lineage and year) for the different coronaviruses compared to reassortment rates in seasonal human influenza A/H3N2 and influenza B viruses as estimated in Müller *et al*. (2020). For OC43 and NL63, the parts of the recombination networks that stretch beyond 1950 are not shown to increase readability of more recent parts of the networks.

The human seasonal coronaviruses all have recombination rates around 1 × 10^−5^ per site and year (Fig. S7). This is around 10 to 20 times lower than the evolutionary rate (Fig. S8). In contrast to the recombination rates, the evolutionary rates vary greatly across the human seasonal coronaviruses, with rates between a median of 1.3 × 10^−4^ (CI 1.1 – 1.5 × 10^−4^) for NL63 and median rate of 2.5 × 10^−4^ (CI 2.2 – 2.7 × 10^−4^) and 2.1 × 10^−4^ (CI 1.9 – 2.3 × 10^−4^) for 229E and OC43 (Fig. S8). These evolutionary rates are substantially lower than those estimated for SARS-CoV-2 (1.1 × 10^−3^ substitutions per site and year (Duchene *et al*., 2020)), which are more in line with our estimates for the evolutionary rates of MERS with a median rate of 6.9 × 10^−4^ (CI 6.0 – 7.9 × 10^−4^). Evolutionary rate estimates can be time dependent, with datasets spanning more time estimating lower rates of evolution that those spanning less time (Duchêne *et al*., 2014). In turn, this means that the evolutionary rates estimates for SARS-CoV-2 will likely be lower the more time passes. It is unclear though, it will approximate the evolutionary rates of other seasonal coronaviruses.

On a per-lineage basis the estimated recombination rate for seasonal coronaviruses translates into around 0.1–0.3 recombination events per lineage and year (Fig. 2E). Recombination events defined here are a product of co-infection, recombination and selection of recombinant viruses. Interestingly, the rate at which recombination events occur is highly similar to the rate at which reassortment events occur in human influenza viruses (Fig. 2D, and Müller *et al*. (2020)). If we assume similar selection pressures for recombinant coronaviruses compared to reassortant influenza viruses, this would indicate similar co-infection rates in influenza and coronaviruses. The incidence of coronaviruses in patients with respiratory illness cases over 12 seasons in western Scotland have been found to be lower (7% – 17%) than for influenza viruses (13%–34% but to be of the same order of magnitude (Nickbakhsh *et al*., 2020). Considering that seasonal coronaviruses typically are less symptomatic than influenza viruses, it is not unreasonable to assume that annual incidence, and therefore likely the annual co-infection rates, are comparable between influenza and coronaviruses.

Compared to human seasonal coronaviruses, recombination occurs around 3 times more often for MERS-CoV (Fig. 2E). MERS-CoV mainly circulates in camels and occasionally spills over into humans (Dudas *et al*., 2018). MERS-CoV infections are highly prevalent in camels, with close to 100% of adult camels showing antibodies against MERS-CoV (Reusken *et al*., 2014). Higher incidence, and thus higher rates of co-infection, could therefore account for higher rates of recombination in MERS-CoV compared to the human seasonal coronaviruses.

The evolutionary purpose of recombination in RNA viruses, as well as whether recombination provides a fitness benefit is unclear (Simon-Loriere and Holmes, 2011). To investigate whether recombination benefits fitness in human coronaviruses, we next tested whether rates of recombination differed on different parts of the genome. To do so, we allowed for different relative rates of recombination within the region 5’ of spike (i.e. mostly ORF1ab), spike itself, and everything 3’ of spike. We computed recombination rate ratios on each of these 3 sections of the genome as the recombination rate on that section divided by the mean rate on the other two sections. We infer that recombination rates are elevated in the spike protein of all human seasonal coronaviruses considered here (Fig. 3, S10 & S11). This is consistent with other work estimating higher rates of recombination on the spike protein of betacoronaviruses (Bobay *et al*., 2020).

**Figure 3:**
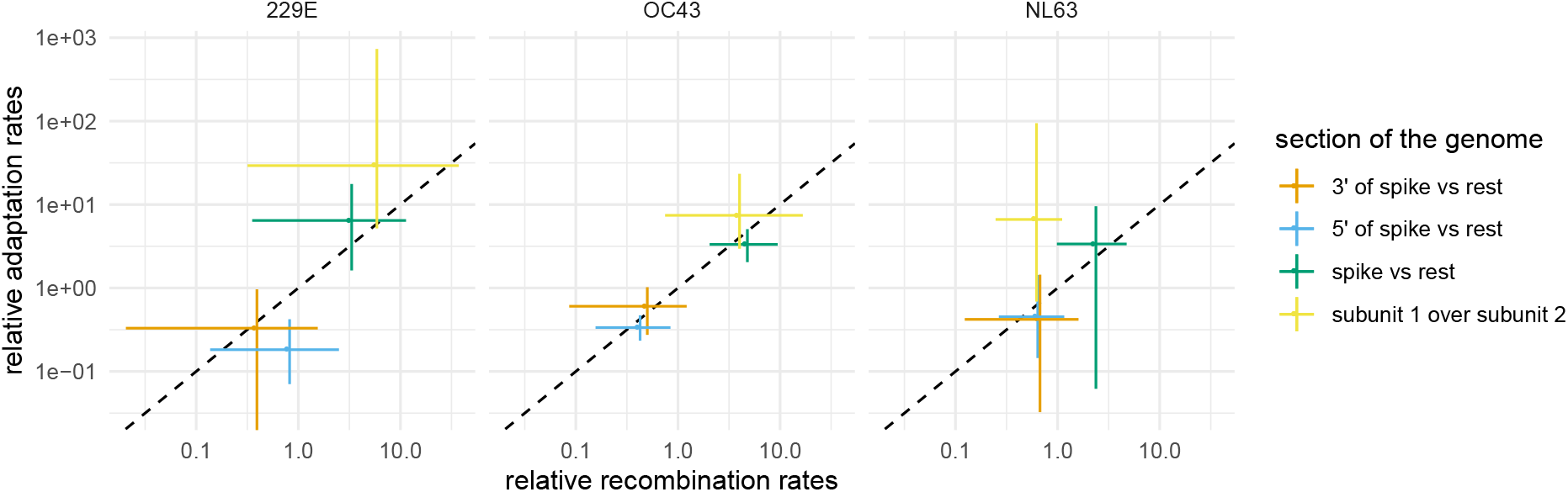
Comparison of recombination rates with rates of adaptation on different parts of the genomes of seasonal human coronaviruses 229E, OC43 and NL63. Relationship between estimated relative recombination rate (x-axis) and relative adaptation rate (y-axis) for three different seasonal human coronaviruses: 229E, OC43 and NL63. These estimates are shown for different parts of the genome, indicated by the different colors. These result from two different types of analysis: one using spike only (subunit 1 over subunit 2, shown in yellow) and one using the full genome (shown in orange, blue and green). The rate ratios denote the rate on a part of the genome divided by the average rate on the two other parts of the genome.

We next tested whether recombination rates are elevated on parts of the genome that also show strong signs of adaptation. To do so, we computed the rates of adaption on different parts of the genomes of the seasonal human coronaviruses using the approach described in (Bhatt *et al*., 2011) and Kistler and Bedford (2021). This approach does not explicitly consider trees to compute the rates of adaptation on different parts of the genomes and is not affected by recombination (Kistler and Bedford, 2021). We find that sections of the genome with relatively higher rates of adaptation correspond to sections of the genome with relatively higher rates of recombination (Fig. 3). In particular, recombination and adaptation are elevated on the section of the genome that codes for the spike protein and are lower elsewhere.

We next investigated whether these trends hold when looking only at spike. The spike protein is made up of two subunits. Subunit 1 (S1) binds to the host cell receptor, while subunit 2 (S2) facilitates fusion of the viral and cellular membrane (Walls *et al*., 2020). S1 contains the receptor binding domain and rates of adaptation have been shown to be high in S1 for 229E and OC43 (Kistler and Bedford, 2021). While the rates of adaptation are relatively low overall for NL63, there is still some evidence that they are elevated in S1 compared to S2 (Kistler and Bedford, 2021).

To test whether recombination rates vary with rates of adaptation on the subunits of spike as well, we inferred the recombination rates from spike only, allowing for different rates of recombination on S1 versus the rest of spike. We find that the rates of recombination are elevated on S1 for 229E and OC43 compared to the rest of the spike gene (Fig. 3). This is consistent with strong absolute rates of adaptation on S1 on these two viruses. For NL63, we find weak evidence for the rate on S2 to be slightly higher than on S1 (Fig. 3), even though the rates of adaptation are inferred to be higher on S1. The absolute rate of adaptation in S1 of NL63 is, however, substantially lower than for 229E or OC43. Additionally, the uncertainty around the estimates on adaption rate ratios between the two subunits for NL63 are rather large and include no difference at all. Overall, these results suggest that recombination events are either positively or negatively selected for. Elevated rates of recombination in areas where adaptation is stronger have been described for other organisms (reviewed here (Nachman, 2002)). Alternatively, higher rates of recombination could also be due to mechanistic reasons, as has been suggested in the case of SARS-CoV-2 (Turakhia *et al*., 2021).

To further investigate this, we next computed the rates of recombination on fitter and less fit parts of the recombination networks of 229E, OC43 and NL63. To do so, we first classify each edge of the inferred posterior distribution of the recombination networks into fit and unfit based on how long a lineage survives into the future. Fit edges are those that have descendants at least one, two, five or ten years into the future and unfit edges those that do not. We then computed the rates of recombination on both types of edges for the entire posterior distribution of networks. Overall, we do not find that fit edges show relatively higher rates of recombination (see figure S9). The simplest explanation is that we do not have enough data points to measure recombination rates on unfit edges, meaning to measure recombination rates on part of the recombination network where selection had too little time to shape which lineages survive and which go extinct. An alternative explanation to why we see elevated rate or recombination in the spike protein, but do not observe a population level fitness benefit could be that most (outside of spike) recombinants could be detrimental to fitness with few (within spike) having little fitness effect at all.

## Conclusion

Though not yet highly prevalent, evidence for recombination in SARS-CoV-2 has started to appear (VanInsberghe *et al*., 2020; Jackson *et al*., 2021; Varabyou *et al*., 2021; Ignatieva *et al*., 2021). As such, it is crucial to know the extent to which recombination is expected to shape SARS-CoV-2 in the coming years, to have methods to identify recombination and to perform phylogenetic reconstruction in the presence of recombination. The results shown here indicate that some recombinants are either positively or negatively selected for. Estimating the deleterious load of viruses before and after recombination using ancestral sequence reconstruction (Yang *et al*., 1995) could help shed light on which sequences are favored during recombination. Furthermore, having additional sequences to reconstruct recombination patterns the seasonal coronaviruses should clarify the role recombination plays in the long term evolution of these viruses.

While their impact on the evolutionary dynamics of SARS-CoV-2 remains unclear, the likely rise of future SARS-CoV-2 recombinants will further necessitate methods that allow phylogenetic and phylodynamic inferences to be performed in the presence of recombination (Neches *et al*., 2020). In absence of that, recombination has to be either ignored, leading to biased phylogenetic and phylodynamic reconstruction (Posada and Crandall, 2002), or non-recombinant parts of the genome have to be used for analyses, reducing the precision of these methods. Our approach addresses this gap by providing a Bayesian framework to infer recombination networks. To facilitate easy adaptation, we implemented the method so that analyses can be set up following the same workflow as regular BEAST2 (Bouckaert *et al*., 2018) analyses. Extending the current suite of population dynamic models, such as birth-death models (Stadler, 2009) or models that account for population structure (Hudson *et al*., 1990; Lemey *et al*., 2009), will further increase the applicability of recombination models to study the spread of pathogens.

## Materials and Methods

### Coalescent with recombination

The coalescent with recombination models a backwards in time coalescent and recombination process (Hudson, 1983). In this process, three different events are possible: sampling, coalescence and recombination. Sampling events happen at predefined points in time. Recombination events happen at a rate proportional to the number of coexisting lineages at any point in time. Recombination events split the path of a lineage in two, with everything on one side of a recombination breakpoint going going in one ancestry direction and everything on the other side of a breakpoint going in the other direction. As shown in Figure 4, the two parent lineages after a recombination event each “carry” a subset of the genome. In reality the viruses corresponding to those two lineages still “carry” the full genome, but only a part of it will have sampled descendants. In other words, only a part of the genome carried by a lineage at any time may impact the genome of a future lineage that is sampled. The probability of actually observing a recombination event on lineage *l* is proportional to how much genetic material that lineage carries. This can be computed as the difference between the last and first nucleotide position that is carried by *l*, which we denote as 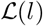. Coalescent events happen between co-existing lineages at a rate proportional to the number of pairs of coexisting lineages at any point in time and inversely proportional to the effective population size. The parent lineage at each coalescent event will “carry” genetic material corresponding to the union of genetic material of the two child lineages.

**Figure 4:**
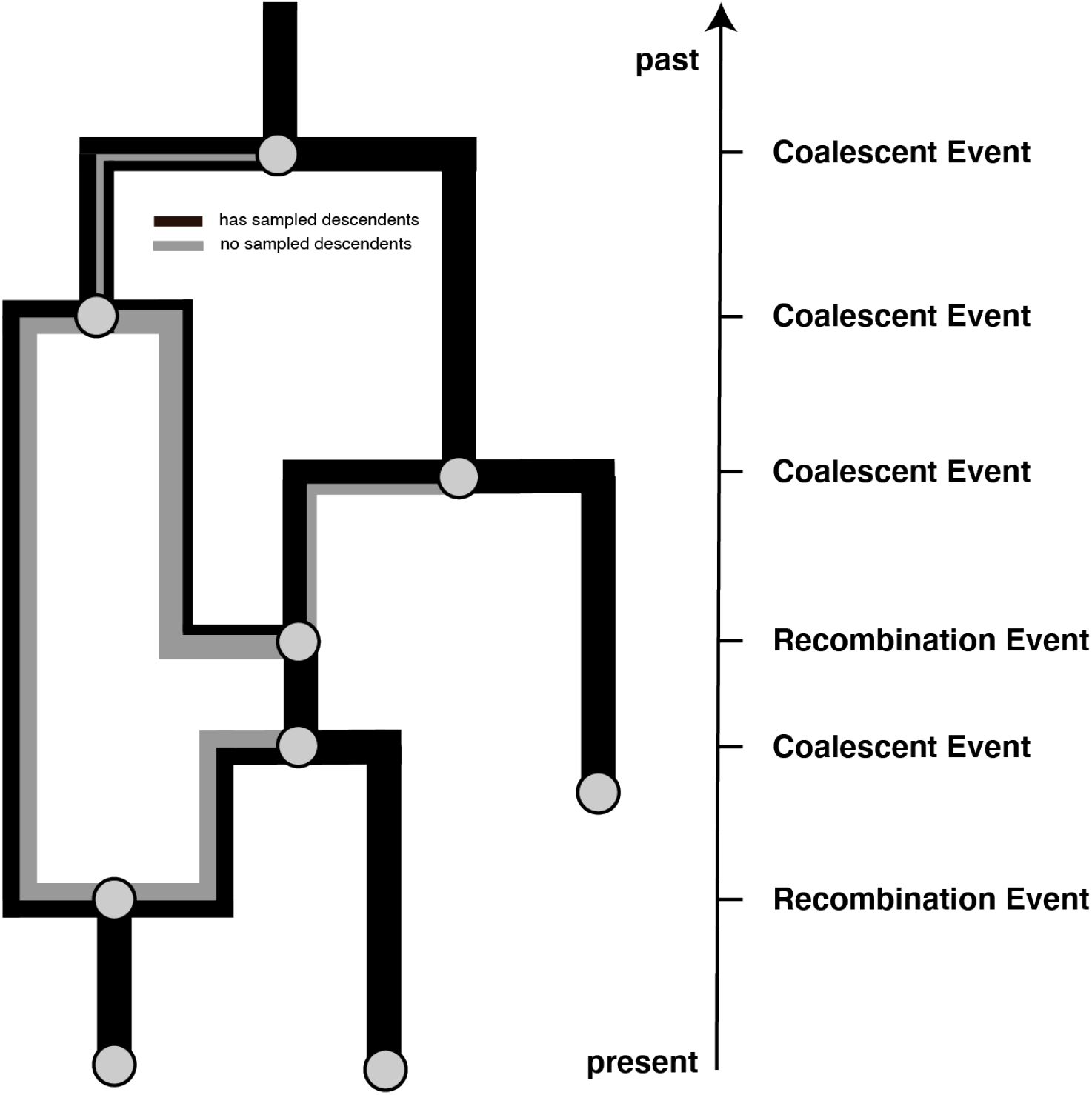
Example recombination network. Events that can occur on a recombination network as considered here. We consider events to occur from present backwards in time to the past (as is the norm when looking at coalescent processes). Lineages can be added upon sampling events, which occur at predefined points in time and are conditioned on. Recombination events split the path of a lineage in two, with everything on one side of a recombination breakpoint going in one direction and everything on the other side of a breakpoint going in the other direction.

### Posterior probability

In order to perform joint Bayesian inference of recombination networks together with the parameters of the associated models, we use a MCMC algorithm to characterize the joint posterior density. The posterior density is denoted as:

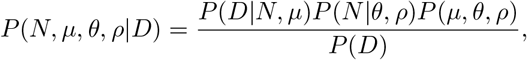

where *N* denotes the recombination network, *μ* the evolutionary model, *θ* the effective population size and *ρ* the recombination rate. The multiple sequence alignment, that is the data, is denoted *D. P*(*D*|*N*, *μ*) denotes the network likelihood, *P*(*N*|*θ*, *ρ*), the network prior and *P*(*μ*, *θ*, *ρ*) the parameter priors. As is usually done in Bayesian phylogenetics, we assume that *P*(*μ*, *θ*, *ρ*) = *P*(*μ*)*P*(*θ*)*P*(*ρ*).

#### Network Likelihood

While the evolutionary history of the entire genome is a network, the evolutionary history of each individual position in the genome can be described as a tree. We can therefore denote the likelihood of observing a sequence alignment (the data denoted *D*) given a network *N* and evolutionary model *μ* as:

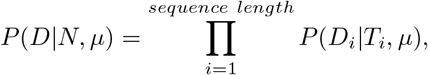

with *D_i_* denoting the nucleotides at position *i* in the sequence alignment and *T_i_* denoting the tree at position *i*. The likelihood at each individual position in the alignment can then be computed using the standard pruning algorithm (Felsenstein, 1981). We implemented the network likelihood calculation *P*(*D_i_*|*T_i_*, *μ*) such that it allows making use of all the standard site models in BEAST2. Currently, we only consider strict clock models and do not allow for rate variations across different branches of the network. This is because the number of edges in the network changes over the course of the MCMC, making relaxed clock models complex to implement. We implemented the network likelihood such that it can make use of caching of intermediate results and use unique patterns in the multiple sequence alignment, similar to what is done for tree likelihood computations.

#### Network Prior

The network prior is denoted by *P*(*N*|*θ*, *ρ*), which is the probability of observing a network and the embedding of segment trees under the coalescent with recombination model, with effective population size *θ* and per-lineage recombination rate *ρ*. It essentially plays the same role that tree prior plays in standard phylodynamic analyses.

We can calculate *P*(*N*|*θ*, *ρ*) by expressing it as the product of exponential waiting times between events (i.e., recombination, coalescent, and sampling events):

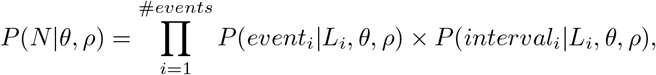

where we define *t_i_* to be the time of the i-th event and *L_i_* to be the set of lineages extant immediately prior to this event. (That is, *L_i_* = *L_t_* for *t* ∈ [*t_i_* – 1, *t_i_*).)

Given the coalescent process is a constant size coalescent and given the i-th event is a coalescent event, the event contribution is denoted as:

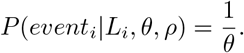

If the i-th event is a recombination event and assuming constant rates of recombination over time, the event contribution is denoted as:

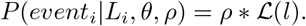

The interval contribution denotes the probability of not observing any event in a given interval. It can be computed as the product of not observing any coalescent, nor any recombination events in interval *i*. We can therefore write:

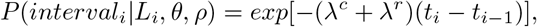

where λ^*c*^ denotes the rate of coalescence and can be expressed as:

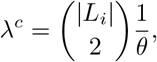

and λ^*r*^ denotes the rate of observing a recombination event on any co-existing lineage and can be expressed as:

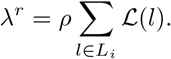

In order to allow for recombination rates to vary across *s* sections 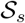 of the genome, we modify λ^*r*^ to differ in each section 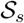, such that:

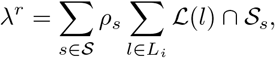

with 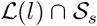 denoting denoting the amount of overlap between 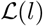 and 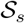. The recombination rate in each section *s* is denoted as *ρ_s_*.

#### MCMC Algorithm for Recombination Networks

In order to explore the posterior space of recombination networks, we implemented a series of MCMC operators. These operators often have analogs in operators used to explore different phylogenetic trees and are similar to the ones used to explore reassortment networks (Müller *et al*., 2020). Here, we briefly summarize each of these operators.

##### Add/remove operator

The add/remove operator adds and removes recombination events. An extension of the subtree prune and regraft move for networks (Bordewich *et al*., 2017) to jointly operate on segment trees as well. We additionally implemented an adapted version to sample re-attachment under a coalescent distribution to increase acceptance probabilities.

##### Loci diversion operator

The loci diversion operator randomly changes the location of recombination breakpoints on a recombination event.

##### Exchange operator

The exchange operator changes the attachment of edges in the network while keeping the network length constant.

##### Subnetwork slide operator

The subnetwork slide operator changes the height of nodes in the network while allowing to change the topology.

##### Scale operator

The scale operator scales the heights of individual nodes or the whole network without changing the network topology.

##### Gibbs operator

The Gibbs operator efficiently samples any part of the network that is older than the root of any segment of the alignment and is thus not informed by any genetic data.

##### Empty loci preoperator

The empty segment preoperator augments the network with edges that do not carry any loci for the duration of a move, to allow larger jumps in network space.

One of the issues when inferring these recombination networks is that the root height can be substantially larger than when not allowing for recombination events. This can cause computational issue when performing inferences. To circumvent this, we truncate the recombination networks by reducing the recombination rate some time after all positions of the sequence alignment have reached their common ancestor height.

#### Validation and testing

We validate the implementation of the coalescent with recombination network prior as well as all operators in the supplement S12. We also show that truncating the recombination networks does not affect the sampling of recombination networks prior to reaching the common common ancestor height of all positions in the sequence alignment.

We then tested whether we are able to infer recombination networks, recombination rates, effective population sizes and evolutionary parameters from simulated data. To do so, we randomly simulated recombination networks under the coalescent with recombination. On top of these, we then simulated multiple sequence alignments. We then re-infer the parameters used to simulate using our MCMC approach. As shown in Figure S13, these parameters are retrieved well from simulated data with little bias and accurate coverage of simulated parameters by credible intervals.

We next tested how well we can retrieve individual recombination events. To do so, we plot the location and timings simulated recombination events for the first 9 out of 100 simulations. We then plot the density of recombination events in the posterior distribution of networks, based on timing and location of the inferred breakpoint on the genome. As shown in figure S14, we are able to retrieve the true (simulated) recombination events well.

We next tested how the speed of inference scales with the number of recombination events, the number of samples in the dataset and the evolutionary rate. To do so, we simulated 300 recombination networks and sequence alignment of length 10,000 under a Jukes Cantor model with between 10 and 200 leafs and a recombination rate between 1 × 10^−5^ and 2 × 10^−5^ recombination events per site per year. This means that for each simulation, there were between 0 and 100 recombination events, allowing us to investigate how the inference scales in different settings. As shown in figure S16, the ESS per hour decreases with the number of recombination events and samples, but not the evolutionary rates. In particular, the ESS per hour decreases much faster with the number of recombination events in a dataset than the number of samples. This suggest that the methods can currently be used more easily to analyze dataset with large number of samples over large number of recombination events.

We next tested how the choice of the prior distribution on the recombination rate impacts the recombination rate estimate. To do so, we simulate 20 recombination networks and sequence alignment of length 10000 under a Jukes Cantor model with 100 leafs and a recombination rate drawn randomly from a log-normal distribution. We then infer the recombination rates using 5 different recombination rate priors as shown in figure S15F that put some or a lot of weight on the wrong parameters. As shown in figures S15A-E, we are able to infer recombination rates, even with the wrong priors.

Additionally, we compared the effective sample size values from MCMC runs inferring recombination networks for the MERS spike protein to treating the evolutionary histories as trees. We find that although the effective sample size values are lower when inferring recombination networks, they are not orders of magnitude lower (see fig S17).

### Recombination network summary

We implemented an algorithm to summarize distributions of recombination networks similar to the maximum clade credibility framework typically used to summarize trees in BEAST (Heled and Bouckaert, 2013). In short, the algorithm summarizes over individual trees at each position in the alignment. To do so, we first compute how often we encountered the same coalescent event at every position in the alignment during the MCMC. We then choose the network that maximizes the clade support over each position as the maximum clade credibility (MCC) network.

The MCC networks are logged in the extended Newick format (Cardona *et al*., 2008) and can be visualized in icytree.org (Vaughan, 2017). We here plotted the MCC networks using an adapted version of baltic (https://github.com/evogytis/baltic).

### Sequence data

The genetic sequence data for OC43, NL63 and 229e were obtained from ViPR (http://www.viprbrc.org) as described in Kistler and Bedford (2021). All virus sequences were isolated from a human host. The sequence data for the MERS analyses were the same a described in Dudas *et al*. (2018), but using a randomly down sampled dataset of 100 sequences. For the SARS like analyses, we used several different deposited SARS-like genomes, mostly originating from bats, as well as humans and one pangolin derived sequence.

### Rates of adaptation

The rates of adaptation were calculated using a modification of the McDonald-Kreitman method, as designed by Bhatt *et al*. (2011), and implemented in Kistler and Bedford (2021). Briefly, for each virus, we aligned the sequence of each gene or genomic region. Then, we split the alignment into 3-year sliding windows, each containing a minimum of 3 sequenced isolates. We used the consensus sequence at the first time point as the outgroup. A comparison of the outgroup to the alignment of each subsequent temporal yielded a measure of synonymous and non-synonymous fixations and polymorphisms at each position in the alignment. This approach requires having sequence data gathered over relatively long time periods where the consensus genome allows for accurate description of the long term evolutionary patterns and, as such, would not be adequate for a pathogen with relatively short evolutionary history, such as for SARS-CoV-2. We used proportional site-counting for these estimations (Bhatt *et al*., 2010). We assumed that selectively neutral sites are all silent mutations as well as replacement polymorphisms occurring at frequencies between 0.15 and 0.75 (Bhatt *et al*., 2011). We identified adaptive substitutions as non-synonymous fixations and high-frequency polymorphisms that exceed the neutral expectation. We then estimated the rate of adaptation (per codon per year) using linear regression of the number of adaptive substitutions inferred at each time point. In order to compute the 5’ of spike and 3’ of spike rates of adaptation we used the weighted average of all coding regions to the left (upstream) or right (downstream) of the spike gene, respectively, using the length of the individual sections as weights. We estimated the uncertainty by running the same analysis on 100 bootstrapped outgroups and alignments.

## Code availability

The Recombination package is implemented as an addon to the Bayesian phylogenetics software platform BEAST2 (Bouckaert *et al*., 2018). All MCMC analyses performed here, were run using adaptive parallel tempering (Müller and Bouckaert, 2020). The source code is available at https://github.com/nicfel/Recombination. We additionally provide a tutorial on how to setup and postprocess an analysis at https://github.com/nicfel/Recombination-Tutorial. The MCC networks are plotted using an adapted version of baltic (https://github.com/evogytis/baltic). All other plots are done in R using ggplot2 (Wickham, 2016) and ggenes (Wilkins, 2019).

## Data availability

The BEAST2 input xml files for all coronavirus analyses in this manuscript, as well as the files used to post process these analyses are available from https://github.com/nicfel/Recombination-Material. The xml files include the sequence data and exact input specification of the coronavirus analyses performed in this manuscript

## Acknowledgments

We would like to thank three anonymous reviewers for their helpful comments that improved the manuscript. We would also like to thanks Timothy G. Vaughan for helpful insights into the implementation of the software. NFM is funded by the Swiss National Science Foundation (P2EZP3_191891). KEK is a NSF GRFP Fellow (DGE-1762114) TB is a Pew Biomedical Scholar and is supported by NIH R35 GM119774. The Scientific Computing Infrastructure at Fred Hutch is supported by NIH ORIP S10OD028685

## Author Contributions

NFM and TB conceived and designed the experiments. NFM and KEK performed statistical analysis and analysed the data. NFM implemented the software. NFM, KEK and TB wrote the paper.

## Supplementary material

**Figure S1:**
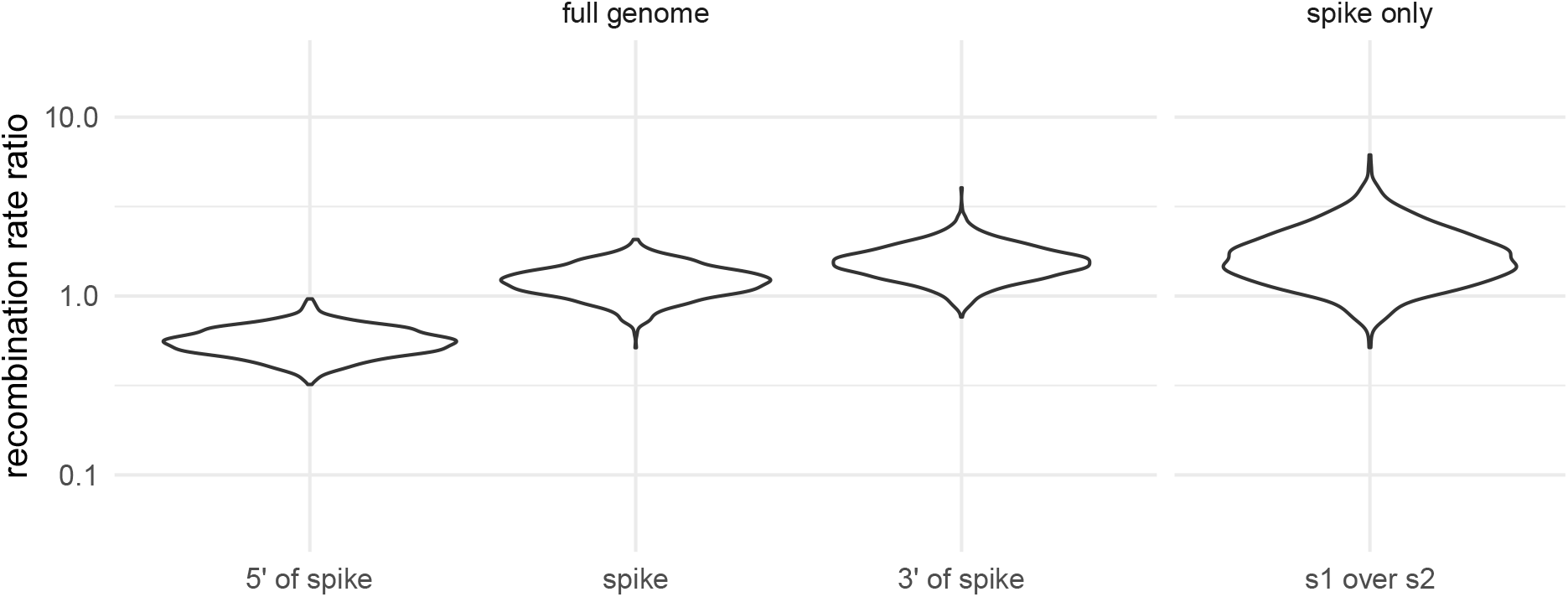
Recombination rate ratios of SARS-like viruses on different parts of the genome. Recombination rate ratios for SARS-like viruses based on two different analyses: one using the full genome (left) and one using the spike protein only (right). The rate ratios denote the rate on a part of the genome divided by the average rate on the two other parts of the genome. s1 over s2 denotes the rate ratio on spike subunit 1 over subunit 2.

**Figure S2:**
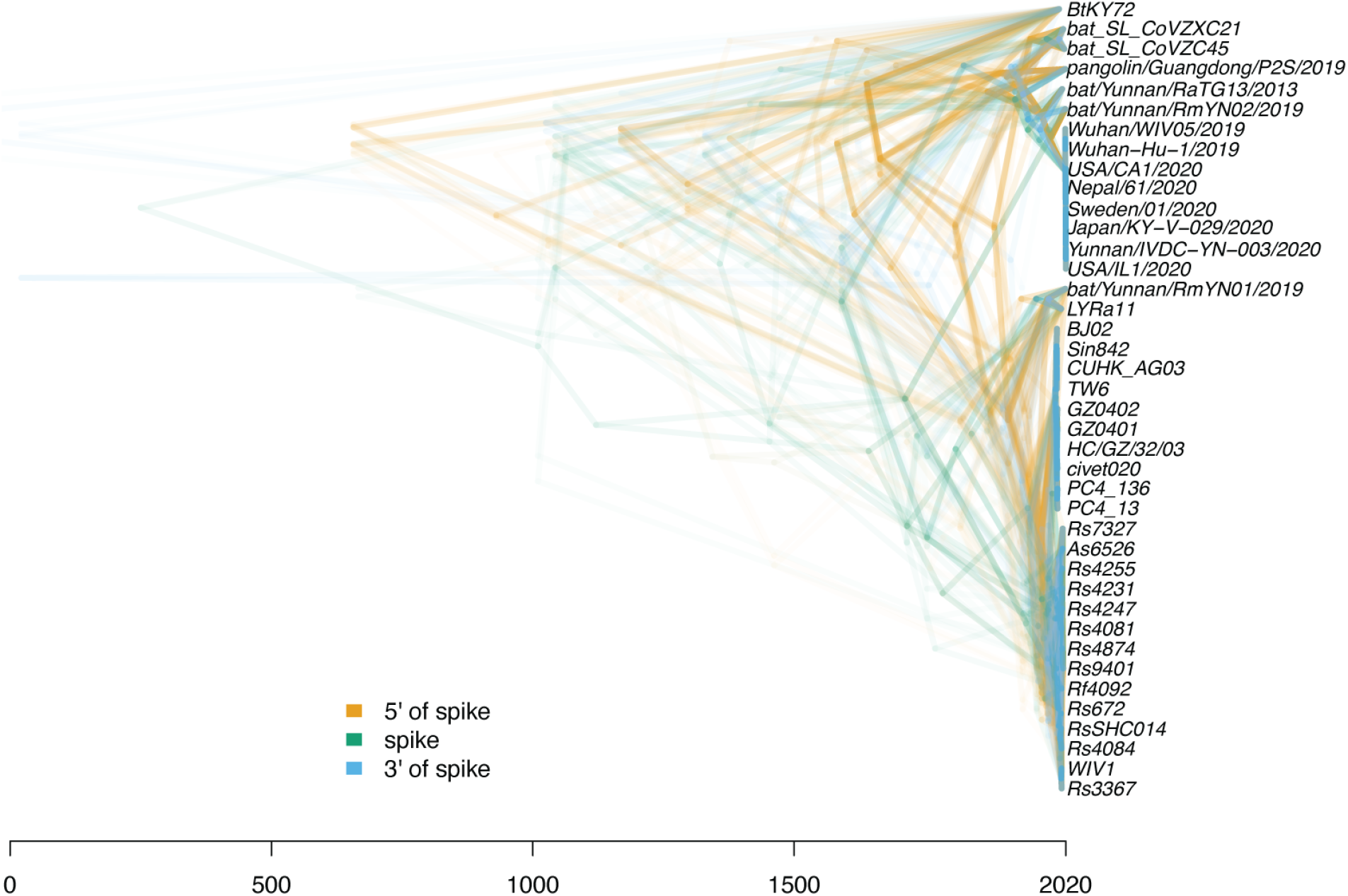
Plot of the local trees of SARS-like virus on different positions across the genome. Densitree (Bouckaert, 2010) plot of local trees in the mcc network of SARS-like viruses. The local trees are shown for every 100th position in the genome and are computed from the mcc network shown in Fig. 1A. The different colors represent whether a local trees was towards the 5’ or 3’ end relative to the region that codes for the spike protein, or whether it was on spike itself.

**Figure S3:**
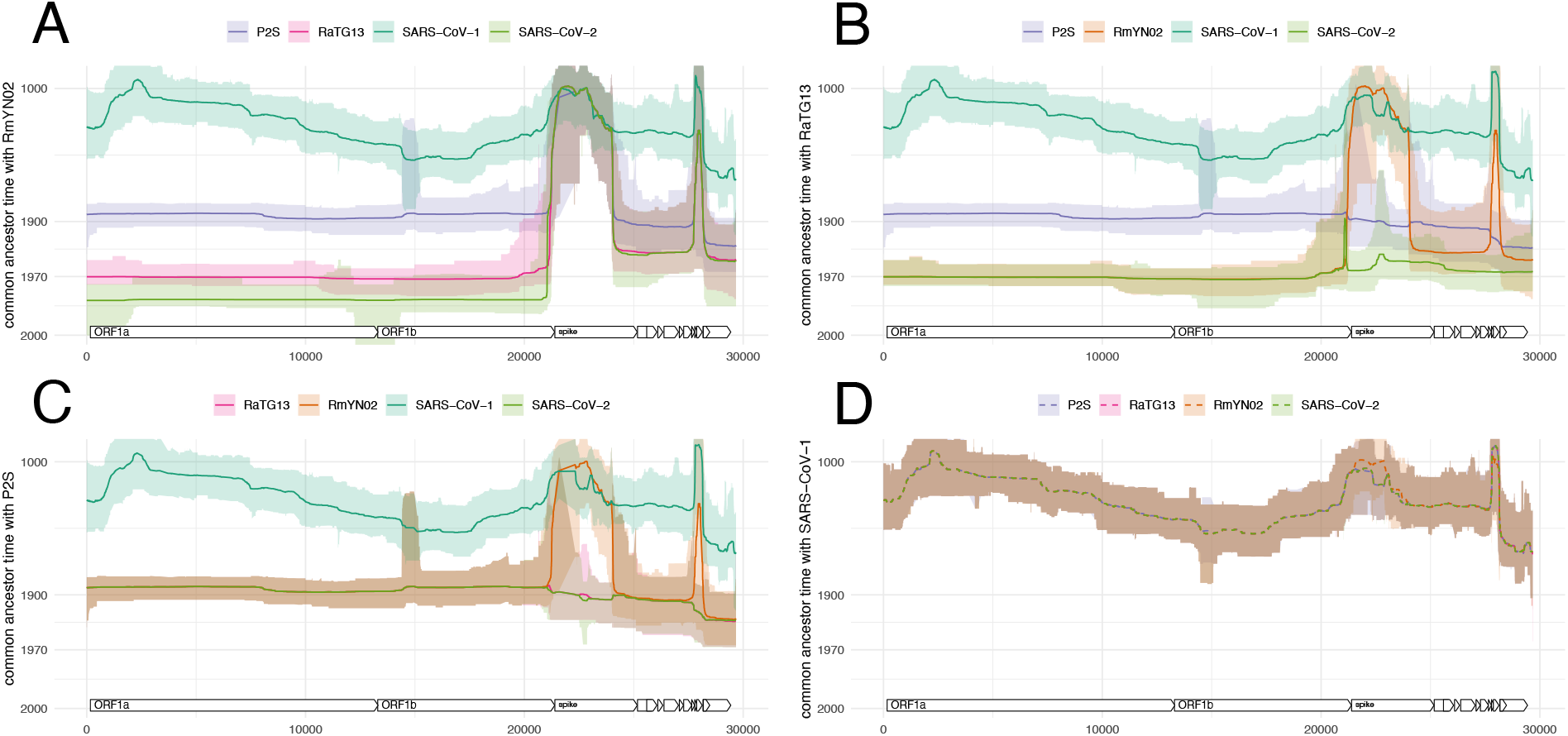
Common ancestor times between sequences of the SARS-CoV-2 clade, as well as SARS-CoV-1. Estimate of common ancestor times of RmYN02 (**A**), RaTG13 **B**, P2S **C** and SARS-CoV-1 **D** with each other and with SARS-CoV-2. The estimates of the common ancestor times assume an evolutionary rate of 5 × 10^−4^. Lower rates would push the common ancestor times further into the past, while higher rates would bring the closer to the present.

**Figure S4:**
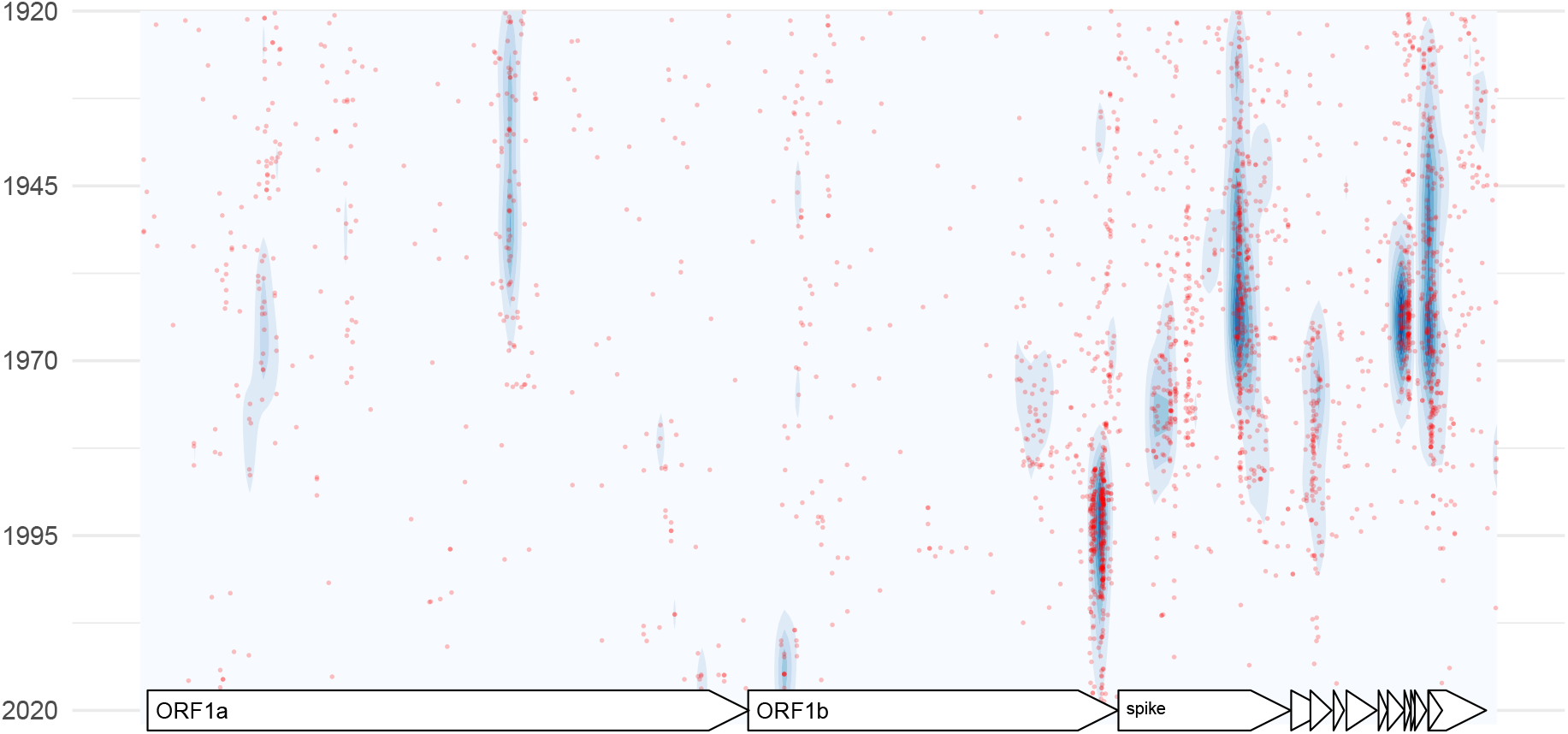
Inferred timings and locations of recombination events ancestral to SARS-CoV-2 in the last one hundred years. Timings and positions of inferred recombination events ancestral to the SARS-CoV-2 lineage are plotted. Each red dot denotes one event in the posterior distribution with the genome position on the x-axis and the year on the y-axis. The density of these events is shown by a contour plot.

**Figure S5:**
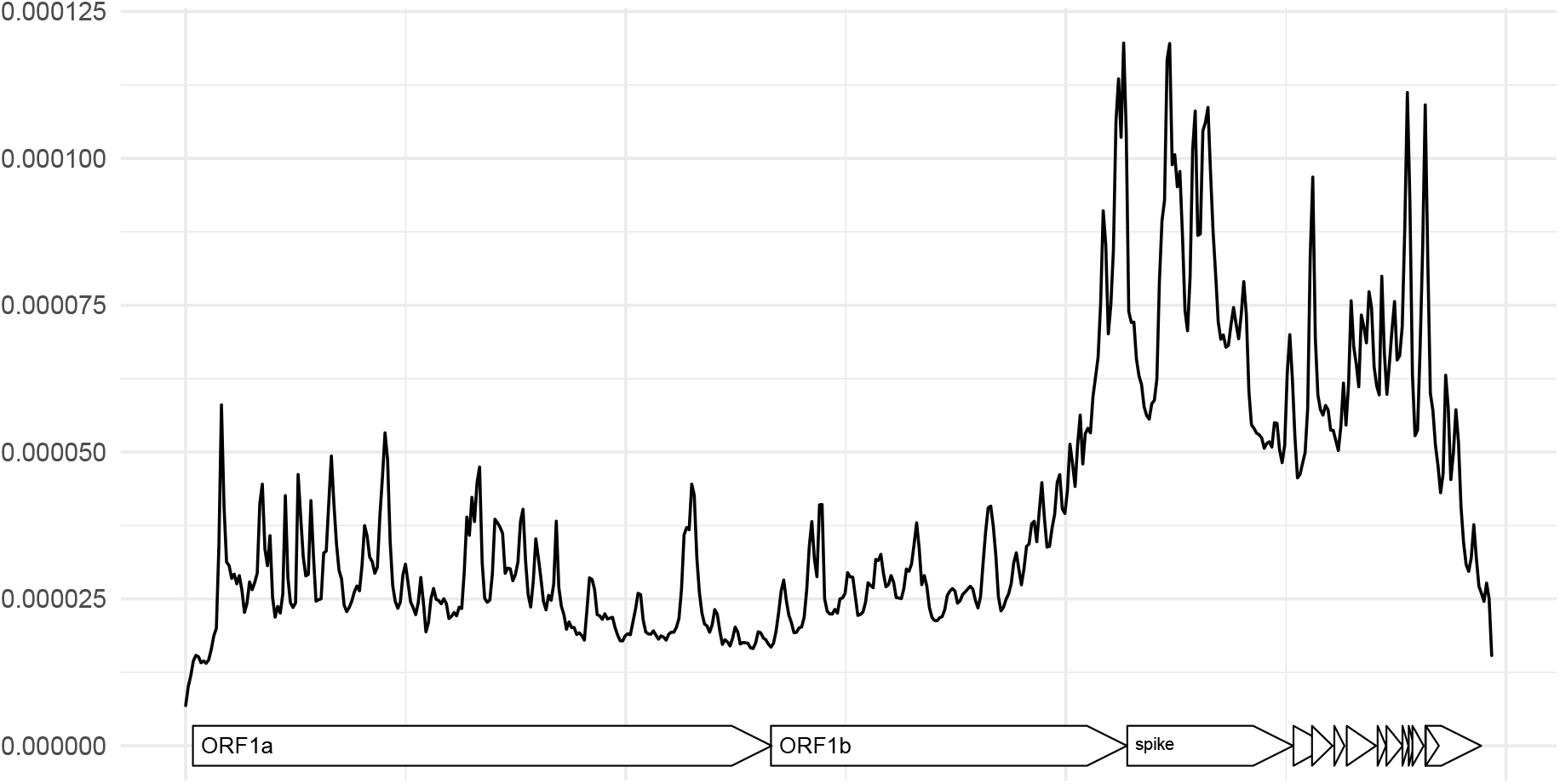
Inferred locations of recombination events in the SARS-like dataset. Here, we show the probability density of recombination events (on the y-axis) along the genome (on the x-axis) of SARS-like viruses.

**Figure S6:**
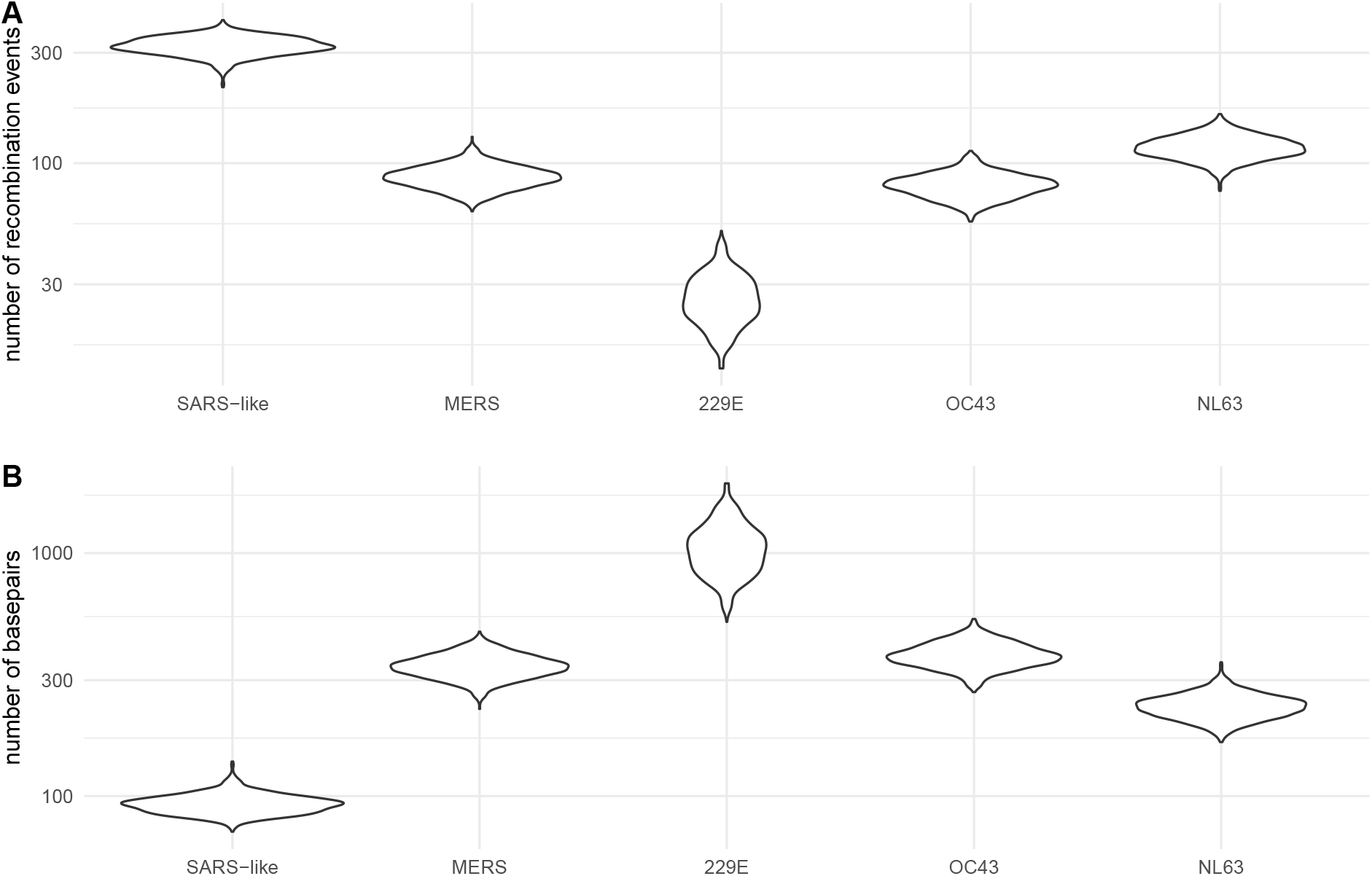
Number of observable recombination events and average length of genomic segment coding for the same tree. **A** Number of recombination events that impact the genome of sampled viruses in the dataset. **B** Average length of a segment in the genome of sampled viruses in the dataset that code for the same phylogenetic tree. That is the average length of a part of the genome that is not broken up by recombination events.

**Figure S7:**
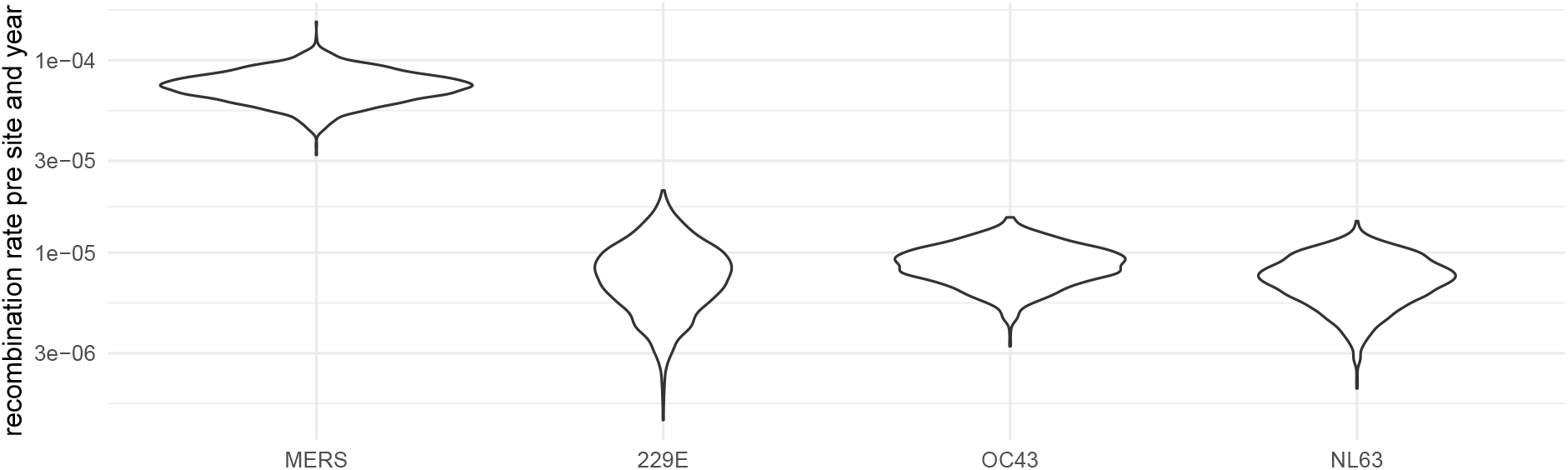
Inferred recombination rates for the different coronaviruses. Posterior distribution of recombination rates per year and per pair of adjacent nucleotides.

**Figure S8:**
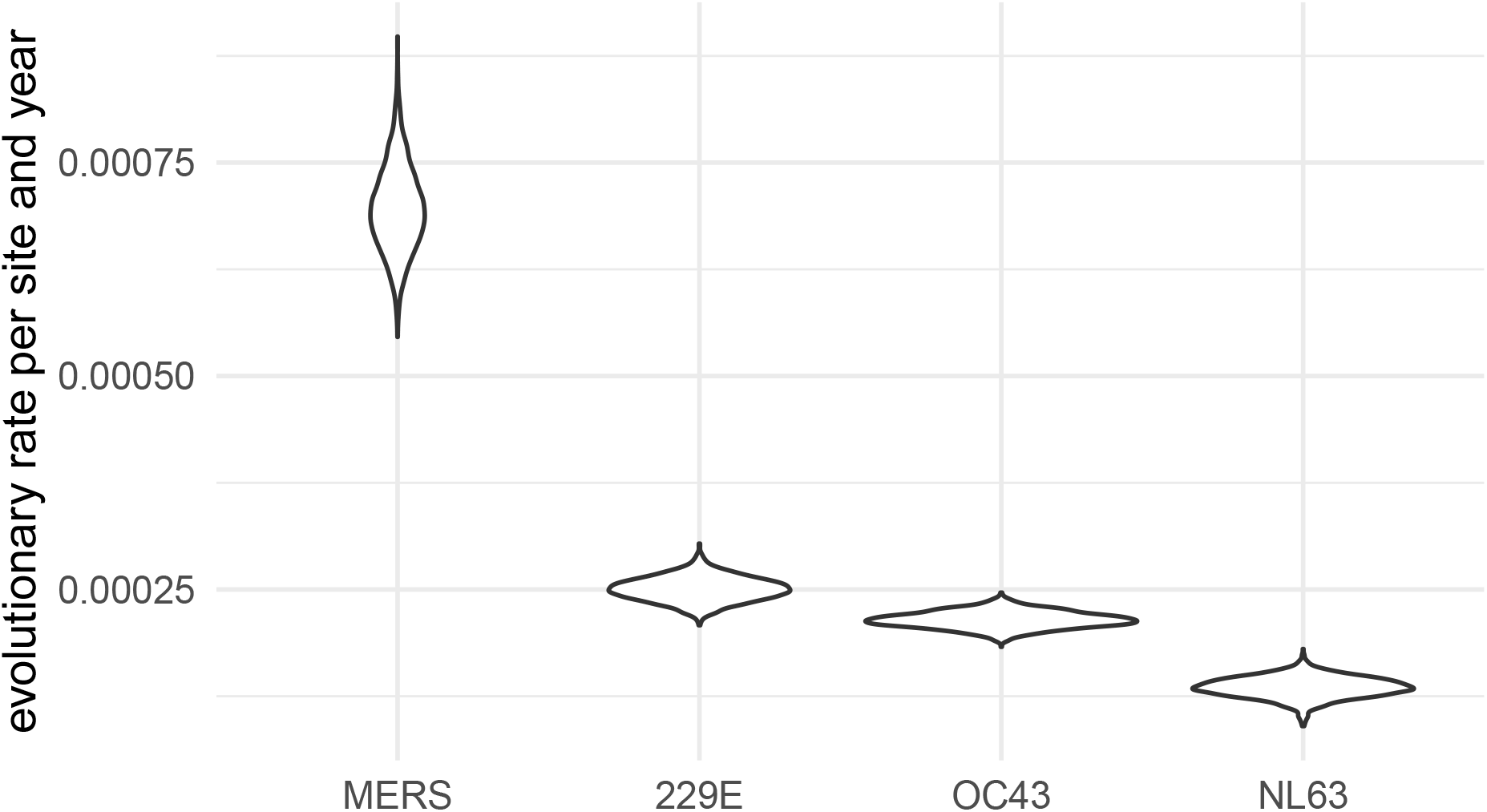
Inferred evolutionary rates for the different coronaviruses. Posterior distribution of evolutionary rates per year and nucleotide.

**Figure S9:**
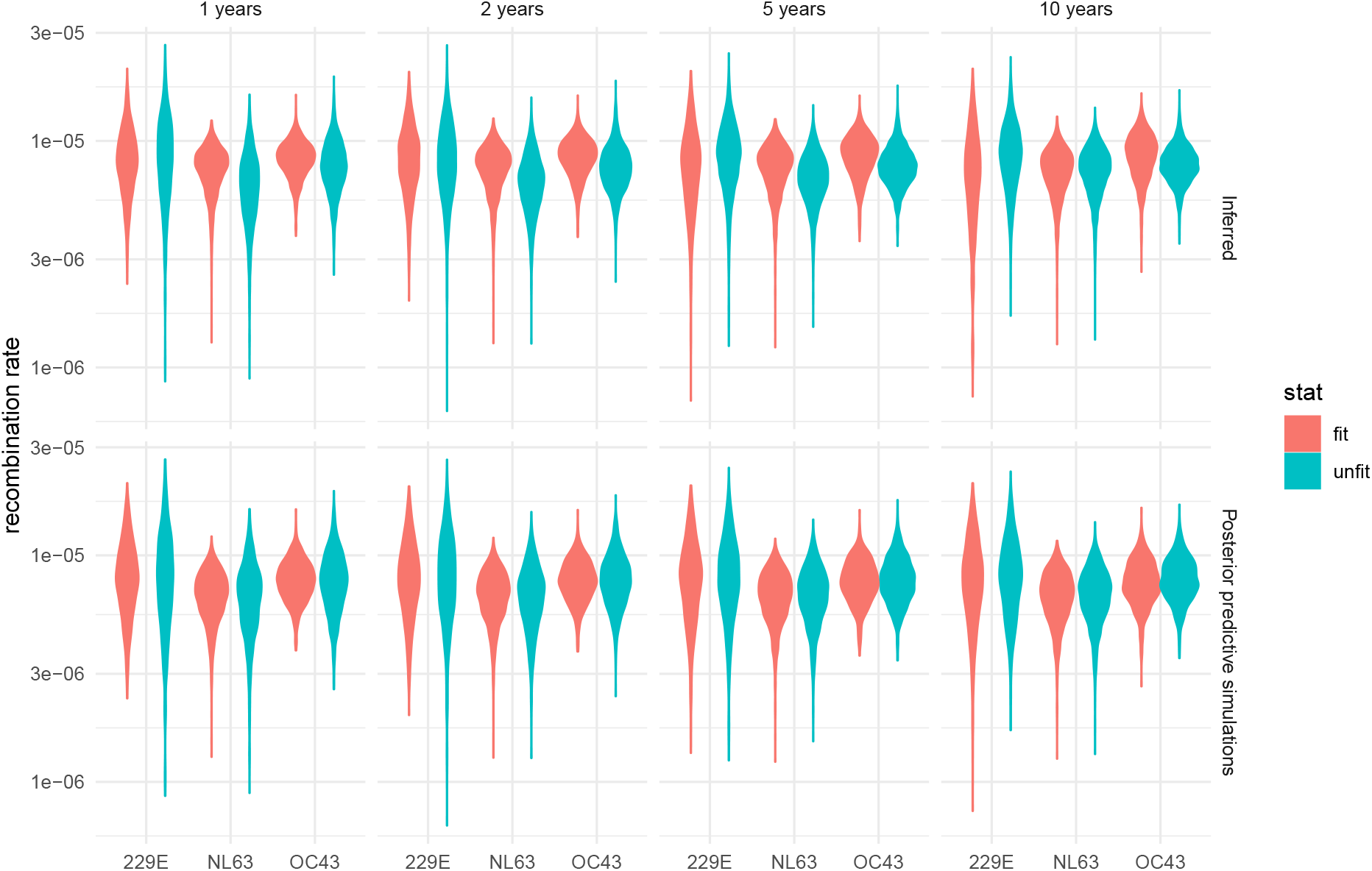
Recombination rates of different parts of the recombination networks. Recombination rates are computed for different parts of the network based on how long lineages persist for into the future. For this analysis, we classified each edge of the recombination network in the posterior distribution into fit and unfit. Fit are edges that persist for at least 1, 2, 5 or 10 years into the future (plots from left to right). We compute the rates of recombination on these edges as well as on those who go extinct more rapidly. We repeat the same for posterior predictive recombination networks that we simulated from the given sampling times, the inferred effective population sizes and the inferred recombination rates under the coalescent with recombination.

**Figure S10:**
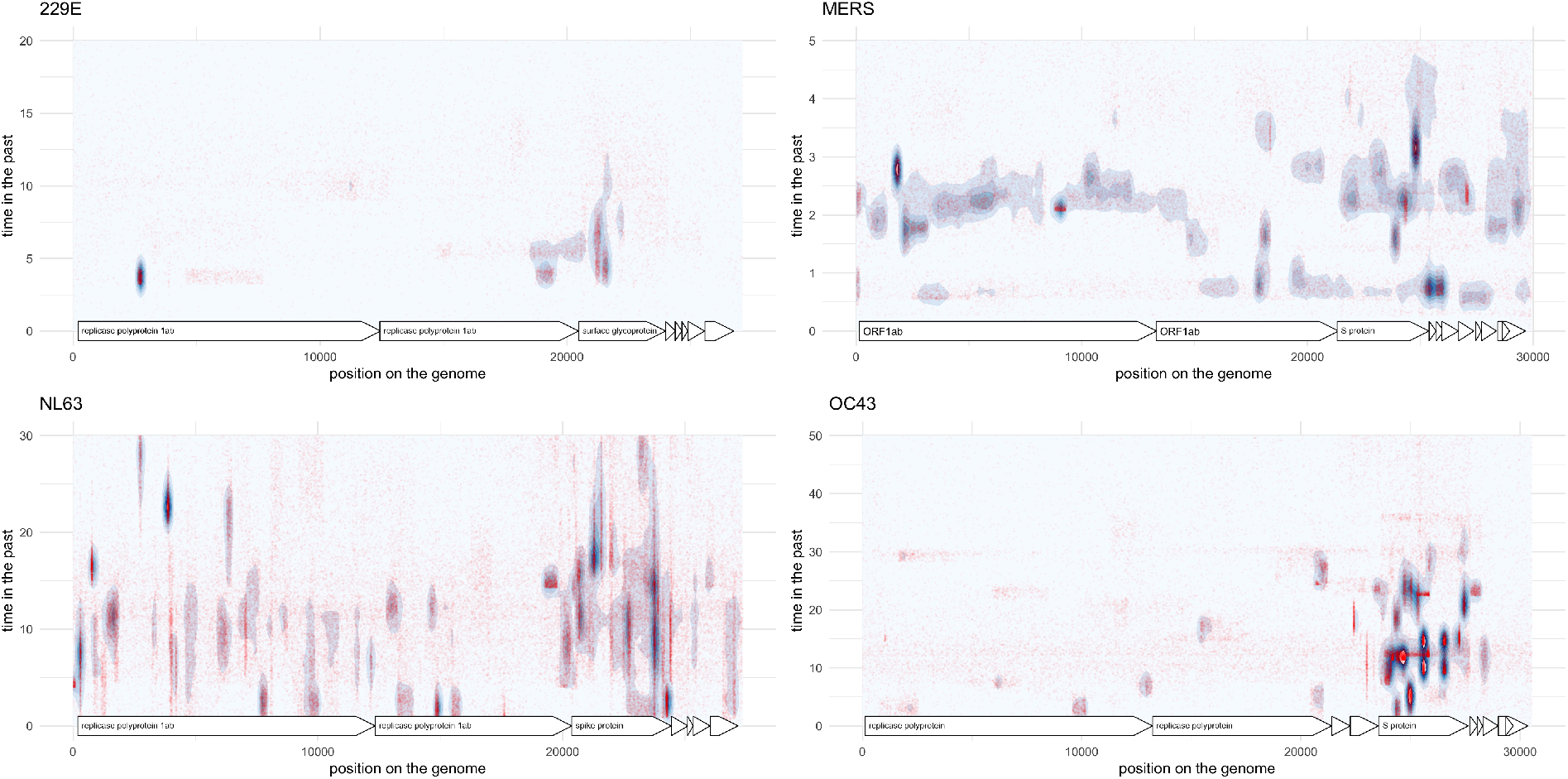
Inferred timings and locations of recombination events of MERS, 229E, OC43 and NL63. Each red dot denotes one event in the posterior distribution with the genome position on the x-axis and the year on the y-axis. The density of these events is shown by a contour plot.

**Figure S11:**
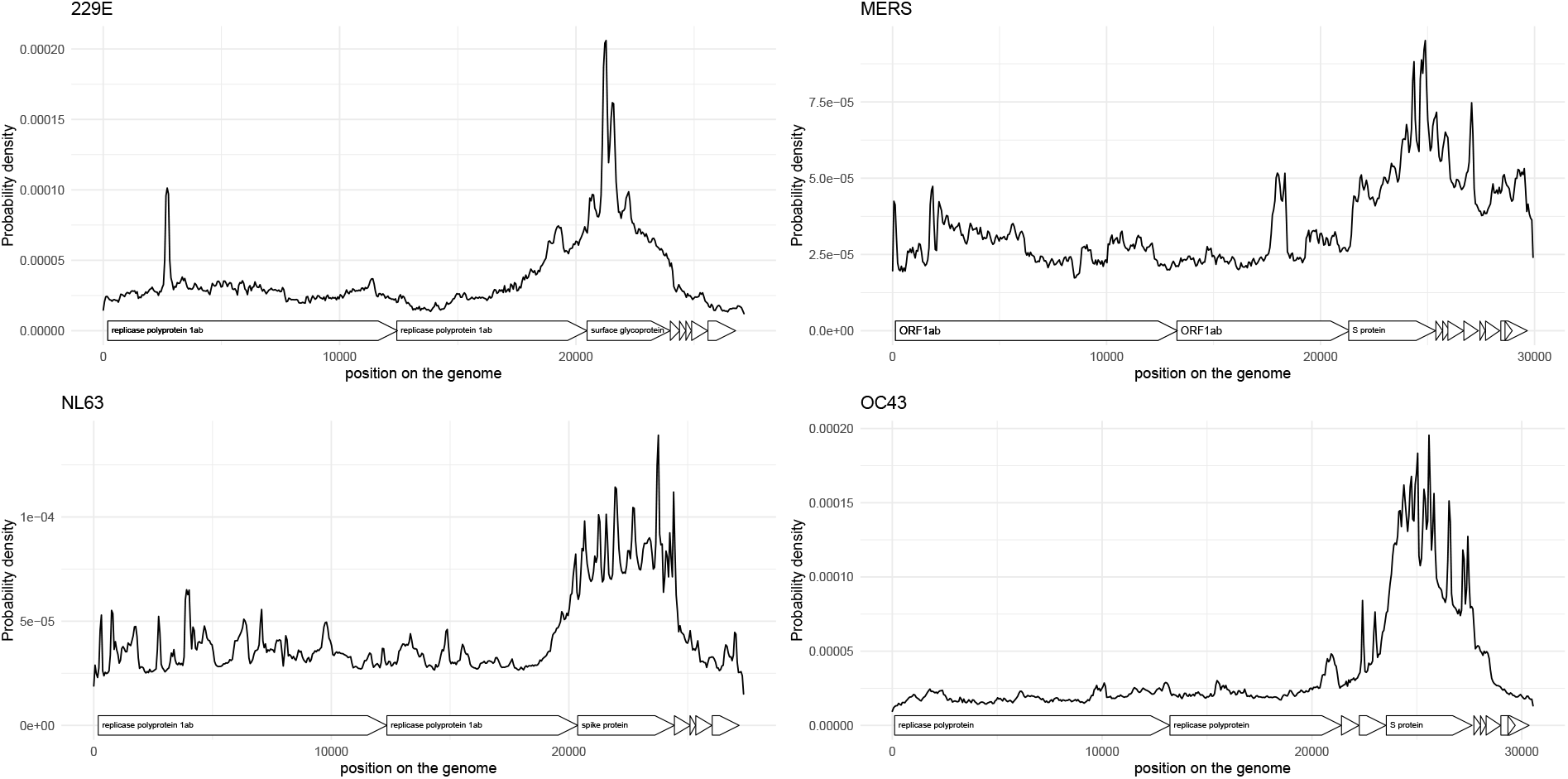
Inferred locations of recombination events of MERS, 229E, OC43 and NL63. Here, we show the probability density of recombination events (on the y-axis) along the genome (on the x-axis) for different coronaviruses.

**Figure S12:**
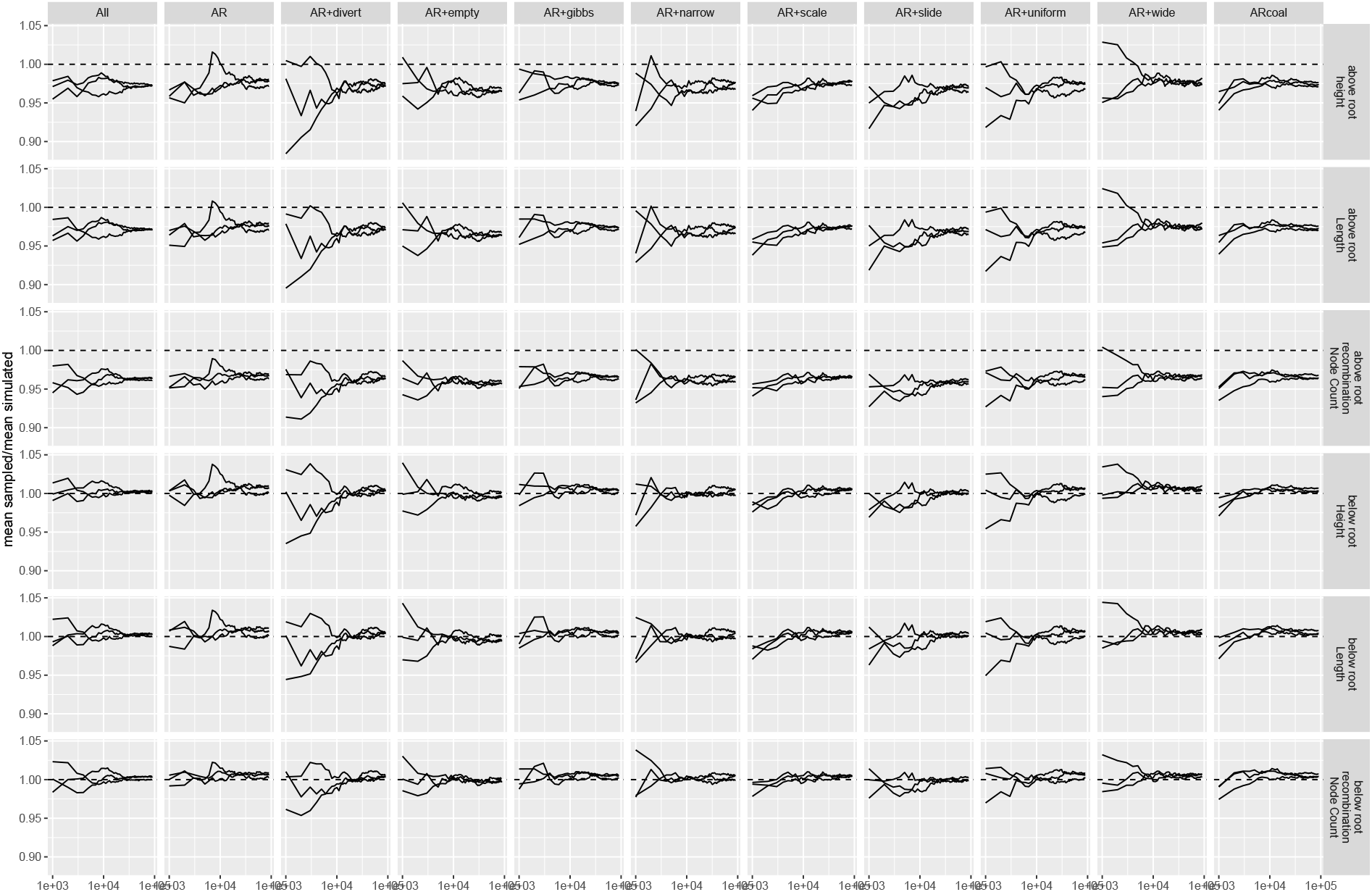
Comparison of network statistics when simulating under the coalescent with recombination compared to sampling under the truncated coalescent with recombination. We compare the posterior distributions of network height, length and the number of recombination nodes when simulating recombination networks under th coalescent with recombination and when MCMC sampling under the implementation of coalescent with recombination. We compare this for all the different MCMC operators implemented. For MCMC operators which are not universal (cannot reach every point in the posterior distribution by themselves), we tested the operator jointly with the Add/remove operator. The statistics “above the root” take into account the full distribution of networks. The statistics “below the root” only take into account the parts of the network that are below (more recent) than the oldest root of any individual position in the alignment. These are the parts of the network that directly impact the likelihood.

**Figure S13:**
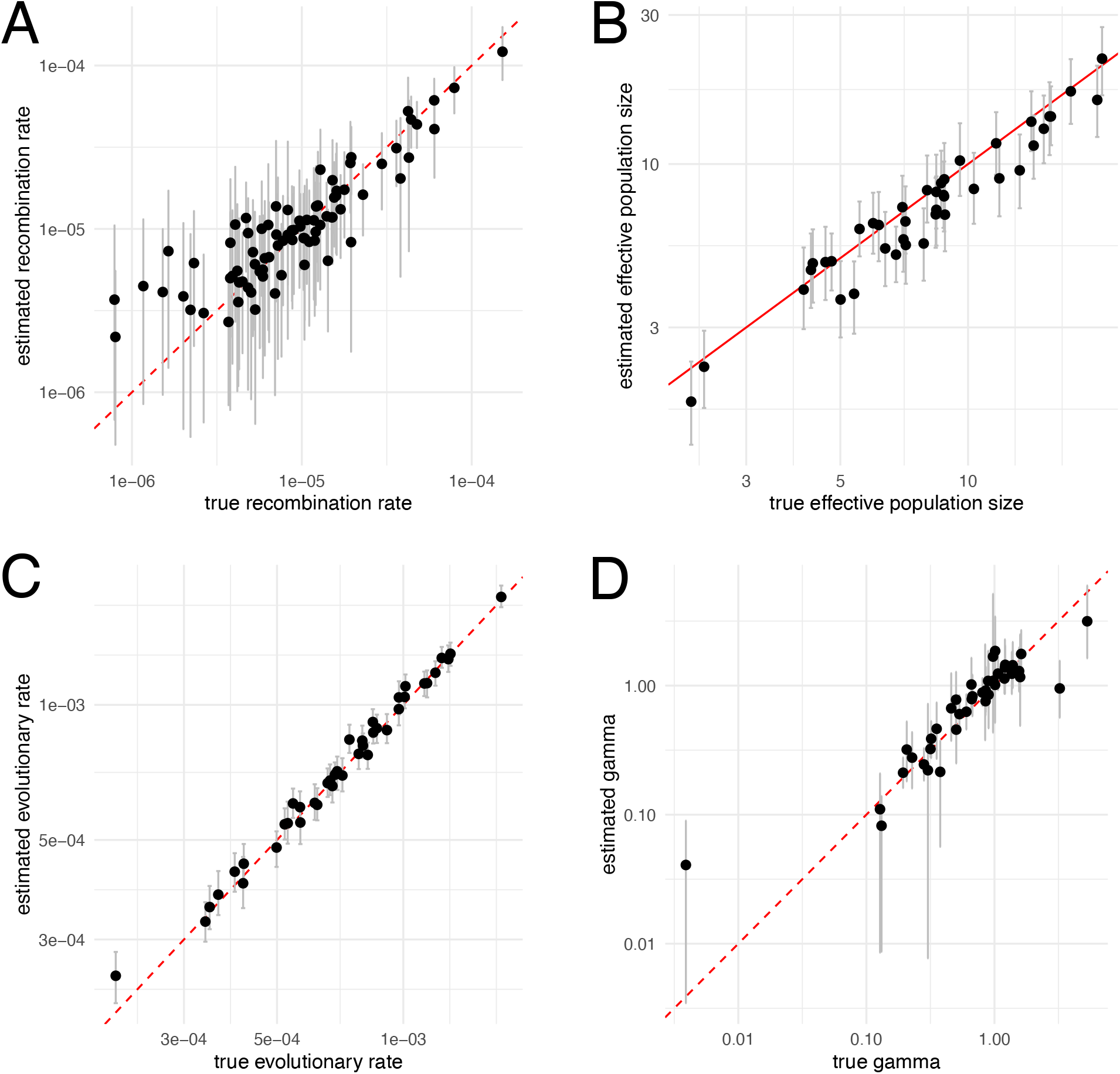
Inferred vs. true rates based on simulated data. We simulated recombination networks and sequence alignment using the randomly drawn values on the x-axis and then re-inferred these parameters on the y-axis. The size of the cross is scaled by the product of the recombination length and the amount of genetic information that recombined. The contour plot shows the distribution of inferred recombination events by location on the genome and time computed from the inferred posterior distribution of networks.

**Figure S14:**
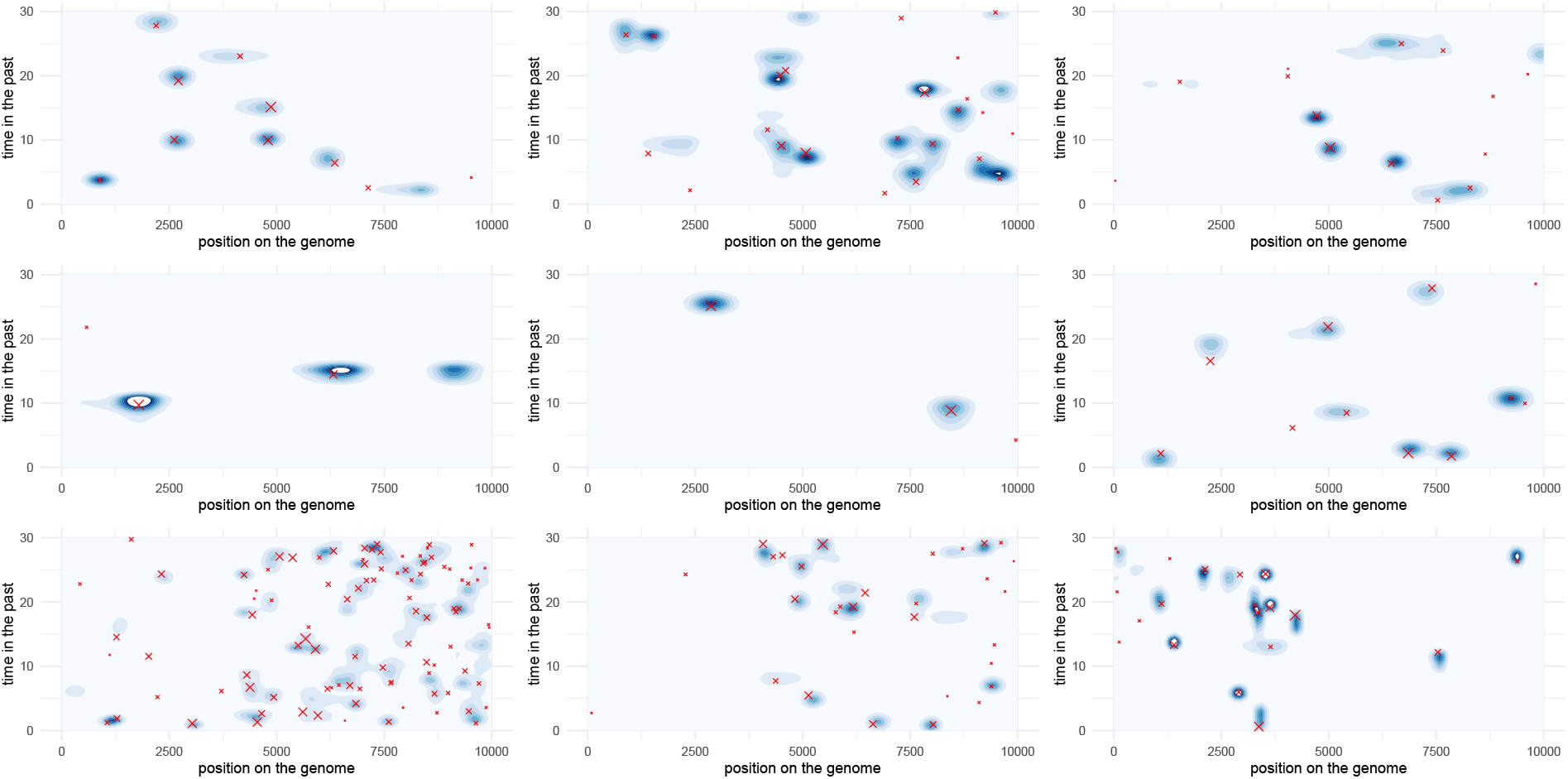
Inferred vs. simulated recombination events. Here, we show the recombination events for the simulated networks (red cross) with the position on the genome on the x-axis and the timing of the event on the y-axis. The contour plots show the density for inferred recombination events for the first 9 iterations of the simulation study. The time in the past is limited to the duration of sampling, i.e. the time when samples were taken.

**Figure S15:**
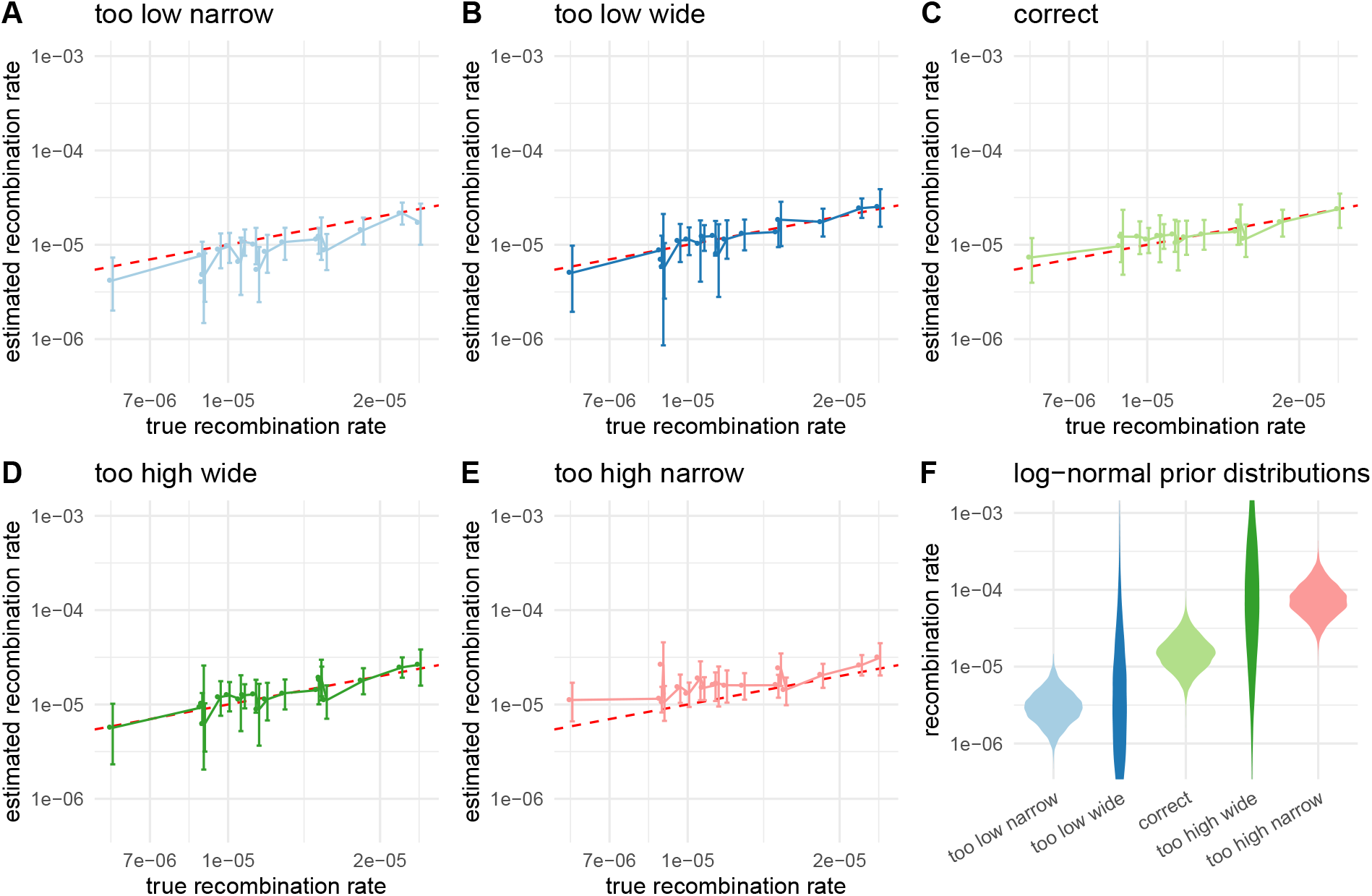
Impact of the recombination rate prior distribution on the inferred recombination rates. Here, we compare then inferred recombination rates when using different prior distributions that differed from the distributions from which the rates for simulations were sampled. The rates for simulations were sampled from a log-normal distribution with *μ* = −11.12 and *σ* = 0.5. In **A**, we shows the inferred rates when using a prior distribution with *μ* = −12.74 and *σ* = 0.5 (leading to a 5 times lower mean in real space than the correct prior). In **B**, we shows the inferred rates when using a prior distribution with *μ* = −12.74 and *σ* = 2. In **C**, we shows the inferred rates when using the same prior distribution as was sampled under. In **D**, we shows the inferred rates when using a prior distribution with *μ* = −9.72 and *σ* = 2. In **E**, we shows the inferred rates when using a prior distribution with *μ* = −9.72 and *σ* = 0.5 (leading to a 5 times higher mean in real space than the correct prior). Figure **F** shows the corresponding density plots for all log normal distributions used as prior distributions on the recombination rates.

**Figure S16:**
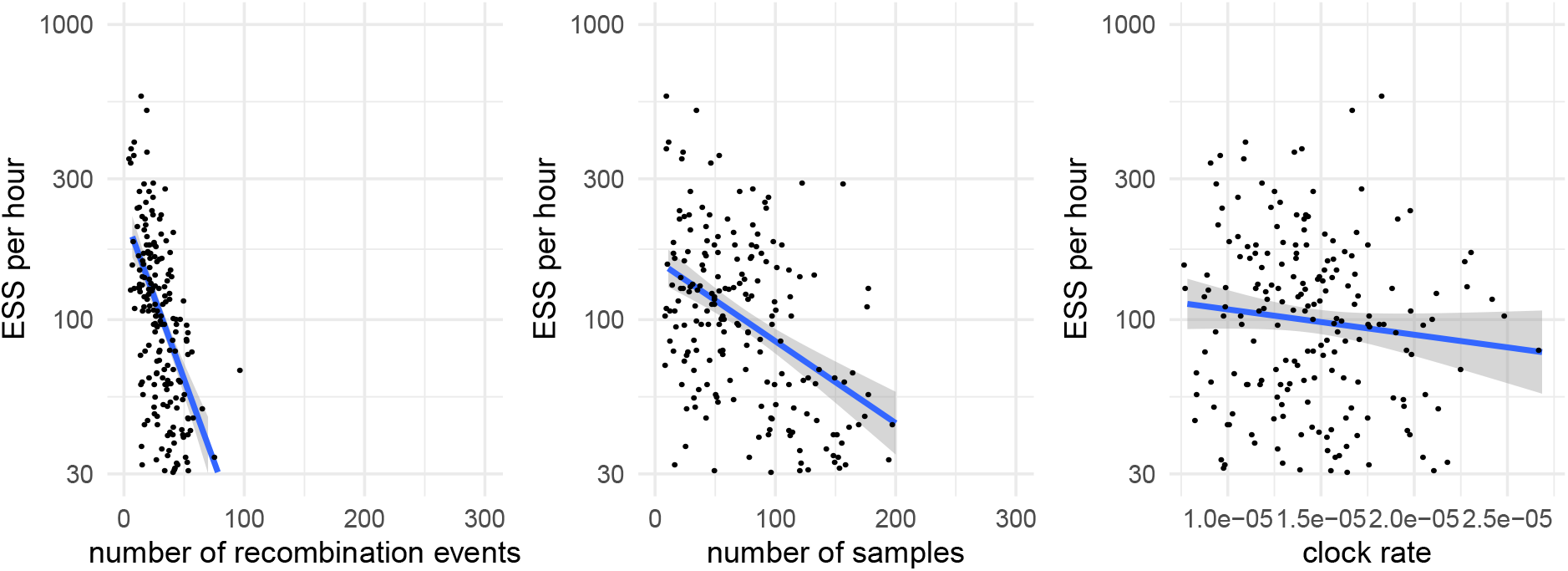
Effective sample sizes per megasample. We computed the effective sample size values computed using coda (Plummer *et al*., 2006) for posterior probabilities, network/tree likelihood values, network/tree root heights and effective population sizes.

**Figure S17:**
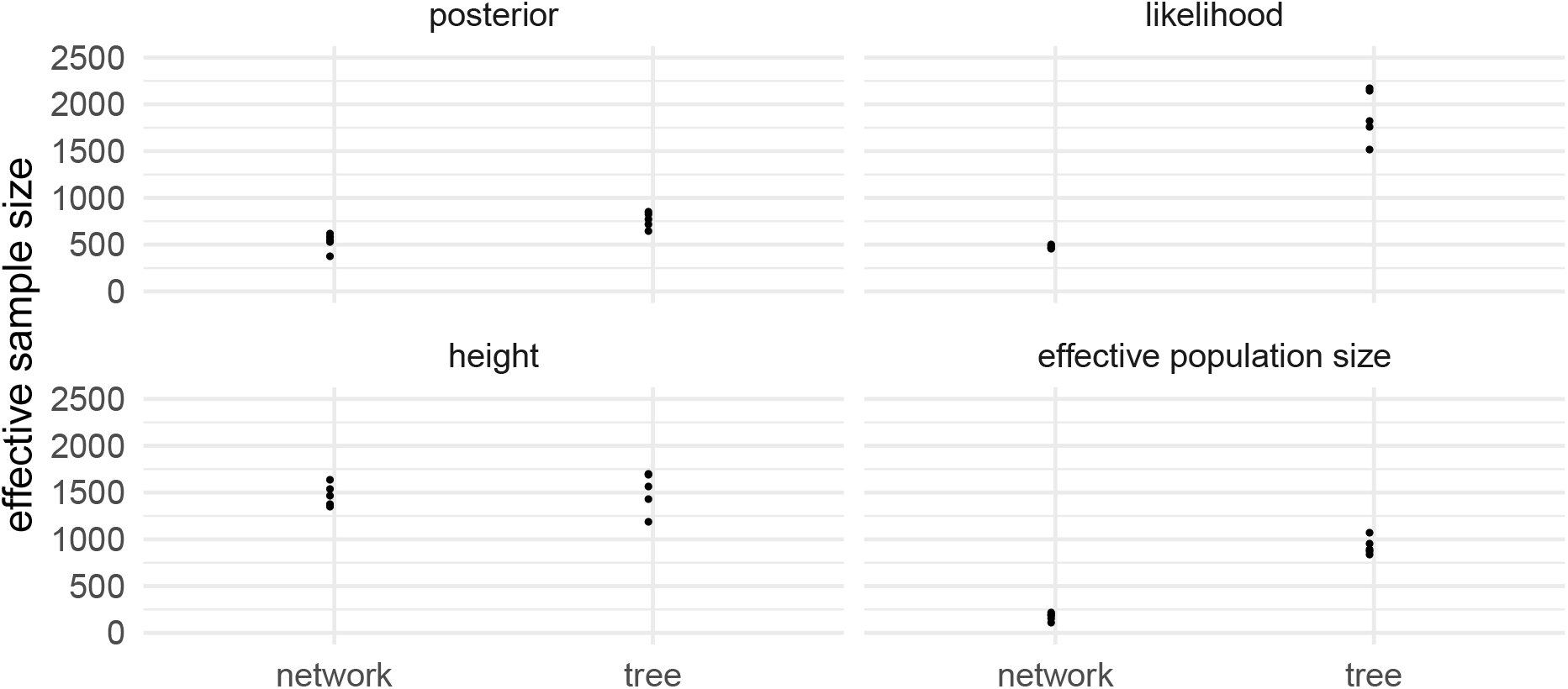
Effective sample Sizes of MERS MCMC runs using the spike protein only. Here, we compare ESS values after 25 Million MCMC iterations when inferring either networks or considering trees only for 100 MERS spike sequences. The operator weights for the inference of recombination networks is the same as used in the other coronaviruses in this manuscript. For the tree inferences, we used the default operator weights. We computed the effective sample size values computed using coda (Plummer *et al*., 2006) for posterior probabilities, network/tree likelihood values, network/tree root heights and effective population sizes.

